# eDNA reveals urban habitat-specific sorting of a mixed regional fish fauna into distinct biodiversity and life-history assemblages

**DOI:** 10.64898/2026.08.29.748007

**Authors:** Katerina L. Zapfe, Elyse Parker, Diego J. Elías, Gabriela M. Hogue, Alex Dornburg

## Abstract

Urbanization is reshaping freshwater ecosystems, with well-documented effects across gradients of land-use change, hydrologic alteration, and habitat degradation. However, how biodiversity is organized among neighboring urban aquatic habitats that differ in hydrologic connectivity, disturbance transmission, residence time, management history, and opportunities for species movement is often less clear. This creates a challenge for interpreting urban fish communities at local scales as species occurrence may reflect both contemporary habitat filtering and historical contingencies including native persistence, interbasin transfer, stocking, and nonindigenous introductions. Here we use eDNA detections, historical records, phylogenetic information, and species trait data to investigate the fish assemblages of the Charlotte metropolitan region. We detect a highly mixed fauna that also depicts a strong signature of structured biodiversity profiles across taxonomic, phylogenetic, functional, and life-history dimensions between habitat types. In particular, bounded habitats contained assemblages with larger-bodied species that are fecund and faster to reproduce relative to free-flowing habitats. Species-level occurrence models did not support a simple trait-by-habitat rule. Instead our results demonstrate that urban aquatic habitats can sort historically mixed regional species pools into predictable assemblage-level life-history profiles while simultaneously retaining signatures of evolutionary and historical biogeographic contingency.

## Introduction

Ecological communities are structured across space and time through interactions among habitat, climate, disturbance, dispersal, and evolutionary history (Leibold et al. 2004; Vellend 2010; Mittelbach and Schemske 2015; Thompson et al. 2020). Broad-scale studies have revealed major patterns in how biodiversity is distributed across landscapes, providing important insight into the environmental processes that shape community composition and species distributions (Newbold et al. 2015; Blowes et al. 2019). However, biodiversity change also unfolds at local scales, where habitat fragmentation, dispersal limitation, fine-grained environmental variation, and local disturbance histories can produce sharp differences among nearby communities (Haddad et al. 2015; Hillebrand et al. 2018; Mori et al. 2018). Under these conditions, broad environmental gradients may give way to finer-scale habitat contrasts, where neighboring systems differ in connectivity, disturbance regime, species composition, and ecological function. Fine-scale studies therefore complement broader environmental analyses by identifying where species, traits, and evolutionary lineages persist, disappear, or are replaced among adjacent habitat types (Magurran et al. 2018).

Ecological disturbance can alter communities simultaneously across taxonomic, functional, and phylogenetic dimensions, often producing uneven patterns of biodiversity change across landscapes (Li et al. 2020; Cancellario et al. 2022). Studies of landscape change have shown that habitat fragmentation, patch configuration, and environmental variation can generate spatial mosaics of filtering, species replacement, and community assembly (Fuller et al. 2015; Reid et al. 2019; Barbarossa et al. 2020). In many systems, these mosaic patterns emerge gradually across broad environmental gradients or habitat transitions (Kraft et al. 2011; Myers et al. 2013). Urbanization can compress these processes into highly fragmented local patchworks, where remnant habitats, engineered systems, altered hydrology, disturbance regimes, species introductions, and built infrastructure occur in close spatial proximity (Alberti et al. 2007; Kondratyeva et al. 2020; Piano et al. 2020). In urban aquatic environments, this compression may be especially pronounced because urban watersheds integrate surrounding land-use change while simultaneously fragmenting habitats through channelization, stormwater infrastructure, impoundment, and altered flow regimes (Walsh and Webb 2016). As a result, closely spaced urban waters may not simply vary in degree of degradation. They may represent different habitat types that transmit, store, and transform disturbance in distinct ways, producing different patterns of species retention, ecological function, and evolutionary diversity (Tickner et al. 2020; Ferzoco and McCauley 2024; Sayer et al. 2025).

Freshwater fish communities are widely used as indicators of ecological condition because assemblage composition responds sensitively to habitat degradation, hydrologic alteration, water quality, land-use change, and biological invasion (Karr 1981; Brown et al. 2009; Wang et al. 2024). However, in urban watersheds, fish assemblages are not shaped by contemporary habitat conditions alone. Species occurrences may also reflect interbasin transfers, sport-fish stocking, management, bait or pet releases, and other forms of human-mediated faunal movement fragmentation that challenge interpretation from species lists (Copp et al. 2005; Leprieur et al. 2008; Kwik et al. 2013; Love et al. 2019). Moreover, freshwater fish assemblages are also inherently difficult to sample consistently across fragmented urban systems. Detectability can vary with habitat structure, sampling gear, water conditions, seasonality, and species behavior that can challenge comparisons across habitats (Fischer and Quist 2014; Dornburg et al. 2017; Mehdi et al. 2021; Bacheler 2026). Environmental DNA (eDNA) sampling offers a complementary approach by providing a standardized snapshot of aquatic community composition across sites previously challenged by logistical constraints to physical sampling (Miya et al. 2015; Shaw et al. 2016; Stoeckle et al. 2017; McElroy et al. 2020). When paired with ecological traits, phylogenetic information, and historical records, eDNA detections can be used to ask whether urban habitat types differ not only in which species are present, but in the functions, lineages, and life-history strategies those species represent.

Freshwater ecosystems are among the most imperiled habitats on the planet (Albert et al. 2021; Hughes 2021) and it is estimated that a third of freshwater fishes face threat of extinction (Hughes 2021; Miranda and Miqueleiz 2021). Thus, the need for an increased understanding of the ecological dynamics in freshwater ecosystems and their biodiversity is particularly acute in urbanized environments. The southeastern United States is a freshwater biodiversity hotspot in North America that contains many endemic taxa, while simultaneously supporting some of the most rapidly expanding urban landscapes in the country (Collen et al. 2014; Elkins et al. 2019; Hilburn et al. 2025; Sayer et al. 2025). Across the region, urban growth is reshaping streams, wetlands, ponds, reservoirs, and tributaries through land-use conversion, hydrologic modification, fragmentation, and infrastructure expansion (Terando et al. 2014; Van Metre et al. 2019; Meador 2020; Nedd and Anandhi 2022). The Charlotte metropolitan region represents an especially pronounced example of these dynamics because it lies within the rapidly expanding urban corridor connecting North Carolina, South Carolina, and Georgia and is projected to continue substantial population and infrastructure growth over the coming decades (Terando et al. 2014; Centralina Regional Council 2020). These conditions make Charlotte a high-stakes test case for examining how contemporary urban freshwater biodiversity is assembled across neighboring aquatic habitats.

Here, we use eDNA-derived fish community data from sites across urban stream networks within the Catawba and Yadkin-Pee Dee urban watersheds in the Charlotte metropolitan region to test whether free-flowing and bounded urban aquatic habitats support distinct environmental, taxonomic, functional, phylogenetic, and life-history assemblages. We first evaluate whether these habitat categories represent distinct environmental contexts. We then quantify species detections and drainage-status categories to determine whether contemporary assemblages reflect a mixed regional fauna shaped by native persistence, interbasin transfer, stocking, pond management, and/or nonindigenous introductions. We next compare biodiversity-retention profiles and species-composition structure to ask whether habitat type organizes taxonomic, phylogenetic, and functional components of diversity. Finally, we examine whether bounded and free-flowing habitats differ in assemblage-level life-history composition, even when individual species occurrence does not follow a simple trait-by-habitat rule. By linking environmental context, faunal mixing, historical records, multidimensional biodiversity, and life-history composition, we show how urban aquatic habitat type can sort a mixed regional species pool into distinct assemblage-level outcomes.

## Methods

### Sample area overview

Fourteen aquatic sites were selected across the Charlotte metropolitan region to capture variation in urban land use and local habitat conditions (**Figure 1**). Candidate sites were identified using the North Carolina Wetland Assessment Map (NC Department of Environmental Quality) and evaluated in Google Earth to achieve a spatially distributed design spanning both interior and exterior portions of the Interstate 485 outer beltway, thereby capturing broad variation in metropolitan development intensity. Sites were chosen to represent wetlands, streams, and ponds while maintaining geographic coverage across the study area and consistency with historic sampling efforts in the region (**Figure 1**; (LeGrand, H., J. Amoroso, and T. Howard. 2026)). As Charlotte is situated at the border between the Catawba and Yadkin-Pee Dee River drainage basins, sites were further selected to ensure representation of both drainage basins (**Figure 1**). Site selection was constrained by the ability to access the water edge, as sampling required transporting equipment on foot from nearby access points. All locations were verified through field reconnaissance to confirm the presence of accessible surface water and suitability for sampling (**Supplemental Fig. S1)**.

**Figure 1.**
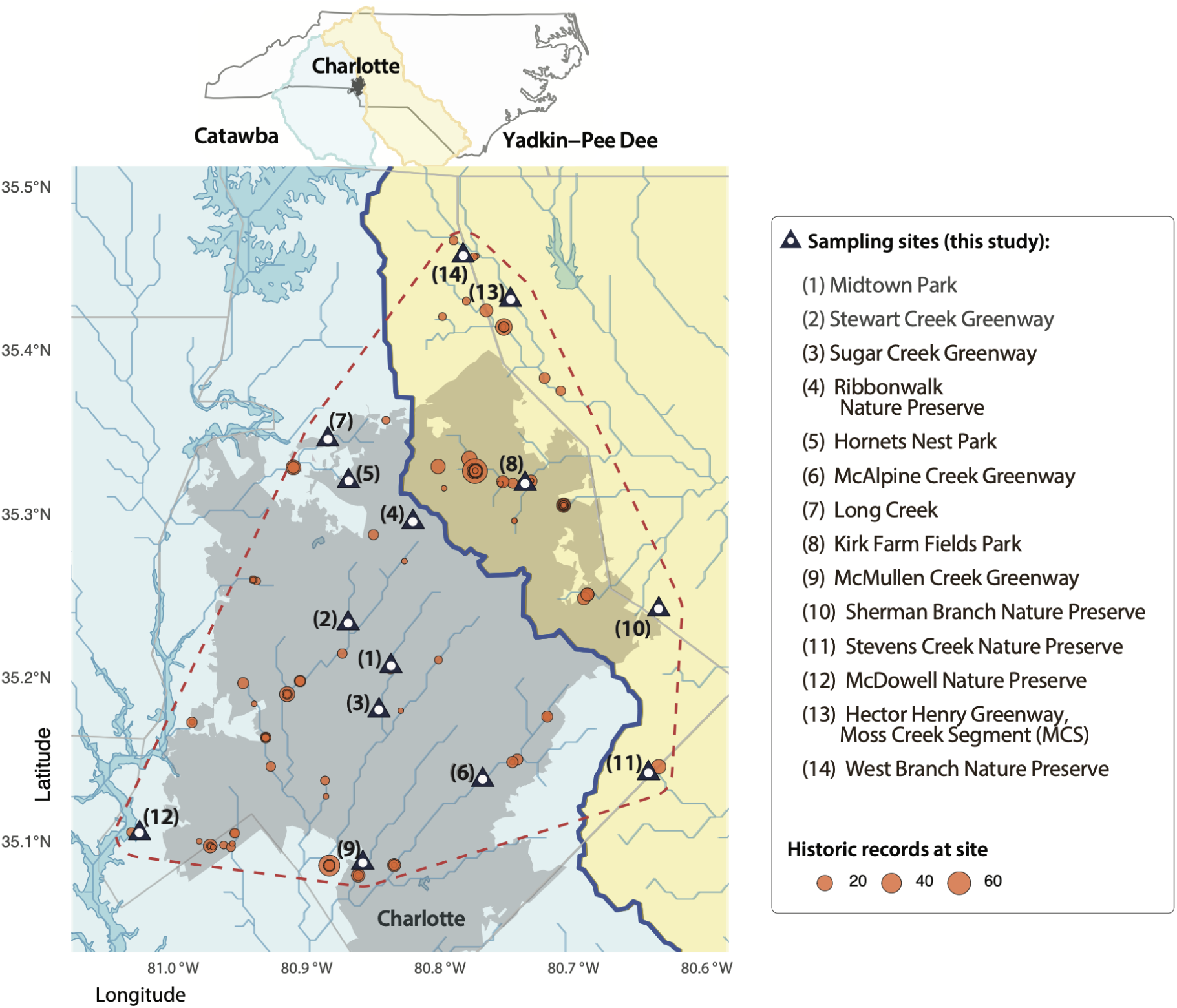
Spatial overview of sample site locations and water drainage network. Sites sampled for this study are indicated by the black triangles and numbered by site name. Historic sampling sites and sample intensities are depicted by circles scaled by sampling intensity. Top insert depicts the placement of Charlotte and highlighted drainage basins within North Carolina. Background shadings delineate the Charlotte city boundary (grey), the Catawba (cool shading), and Yadkin-Pee Dee (warm shading) drainage basins. The solid line delineates the drainage boundary in the Charlotte region. Dotted polygon indicates the extent of our target sampling area. Additional photographs of sites are available in **Supplemental Figure S1**.

**Figure 2.**
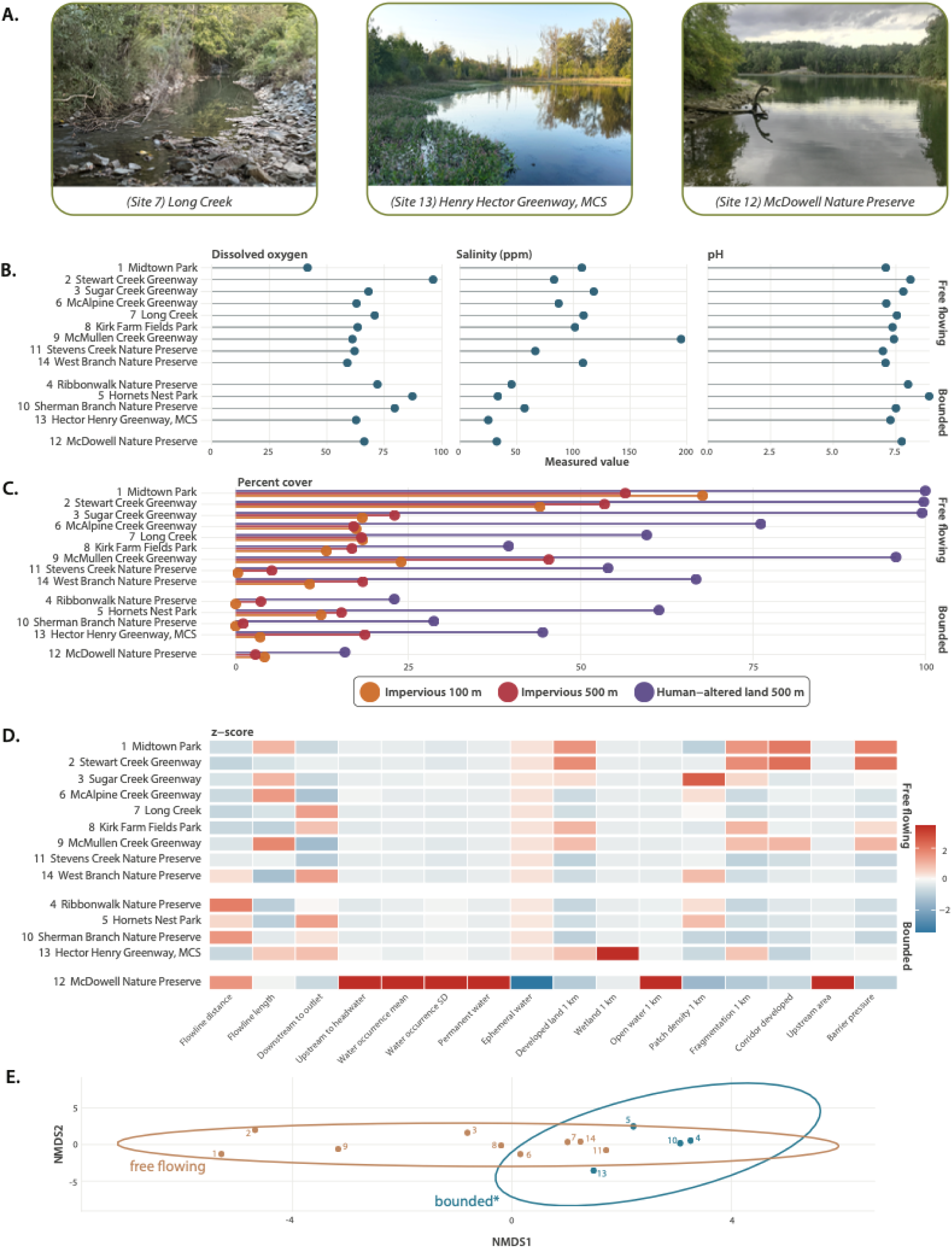
Environmental profiles of sampling sites. (A) Representative photos of sites within each environmental classification category. (B) Dissolved oxygen, salinity, and pH levels (X-axis) between sites (Y-axis). (D) Z-scores of all environmental and GIS data between sites. (E) Environmental NMDS based on z-scores of measured and GIS variables. Points represent sites and shadings indicate habitat type with convex hulls indicating the extent of habitat-type grouping. *indicates the exclusion of the bounded large site due to its high environmental dissimilarity (See supplemental figure S2). Sites in B-D are grouped by habitat type and ordered by distance from the Charlotte city center within groups. Distance from the urban core was calculated as the Euclidean distance from each sampling site to the intersection of Trade and Tryon Streets in Uptown Charlotte (35.2271°, −80.8431°), which represents the historic center of Charlotte’s urban development (Charlotte 2026).

### Landscape alteration metrics

To quantify the degree of landscape alteration around our focal urban aquatic sites, we compiled a suite of GIS-derived metrics capturing variation in hydrologic connectivity, landscape fragmentation, and anthropogenic disturbance using publicly available spatial datasets. Site coordinates were used to define a Charlotte-area spatial extent with a 0.2° buffer, and all vector and raster layers were clipped to this extent prior to analysis. Spatial analyses were conducted after projecting site points and vector layers to UTM Zone 17N (EPSG:32617) to preserve local distance, area, and buffer calculations. We first computed the distance from each sampling site to the nearest mapped river or stream reach and extracted reach length, downstream distance to outlet, upstream distance to headwaters, and upstream contributing area using the HydroSHEDS, HydroRIVERS v1.0, and HydroBASINS level 12 datasets (Lehner and Grill 2013) For calculation of each variable, each site was assigned to the nearest HydroRIVERS flowline, and HydroBASINS level 12 polygons were used to provide watershed context and fallback upstream-area estimates where HydroRIVERS values were unavailable. Next, four metrics capturing surface-water persistence around each site were quantified from the Joint Research Center Global Surface Water occurrence database (Pekel et al. 2016). Within 1 km buffers around each site, we calculated mean occurrence (the average percentage of time that water was present across pixels within the buffer), standard deviation of occurrence (the variability of water persistence within the buffer), the fraction of pixels with occurrence ≥80% (used as a proxy for persistent occurrence of surface water), and the fraction of pixels with occurrence ≤20% (a proxy for rare or intermittent surface-water occurrence). Lastly, to characterize the land-cover context surrounding each site, we quantified the proportion of land classified as built-up (indicating urban development), wetland (signifying semi-aquatic habitat and potential hydrologic connectivity), or open water (representing nearby aquatic habitat availability) within 1 km buffers using ESA WorldCover 2021 v200 (Zanaga et al. 2022)

In addition to the variables representing hydrologic connectivity, we calculated several GIS-derived custom indices that were used as proxies for landscape fragmentation and potential barriers to hydrologic exchange. Developed-patch density was calculated as the number of 4-neighbor connected developed-pixel patches from the ESA WorldCover 2021 land cover raster divided by the total raster-cell count within each 1 km buffer, and a local fragmentation index was calculated as the mean of developed fraction and developed-patch density. Corridor development was estimated as the fraction of developed land within a 120 m buffer surrounding the shortest line connecting each site to its nearest HydroRIVERS flowline to quantify immediate development pressure along the local hydrologic linkage rather than the broader surrounding landscape. Finally, a barrier-pressure index was calculated as the mean of local developed fraction and corridor developed fraction.

Anthropogenic disturbance surrounding each sampling site was quantified using metrics that capture multiple different sources of human-altered land use. We first assessed impervious surface cover, a widely-used proxy for hydrologic and chemical disturbance associated with increased stormwater runoff, altered flow regimes, elevated sediment and contaminant delivery, thermal pollution following precipitation events, and reduced riparian infiltration capacity (Roy et al. 2003; Schiff and Benoit 2007; Dugan et al. 2017a). Imperviousness was calculated as the area-weighted mean percent impervious surface around each sampling site using a 2024 fractional impervious surface raster at 30 m resolution, where each pixel represented the proportion of land covered by impervious materials. A 100 m buffer around each sampling site was used to compute local imperviousness, which represents fine-scale alteration in the immediate riparian or shoreline environment, such as nearby roads, parking lots, culverts, bank hardening, and shoreline modification, which can influence aquatic habitat quality independently of broader watershed context (Roy et al. 2003; Alberts et al. 2018). Imperviousness was also quantified at a landscape-scale within a 500 m buffer to summarize broader surrounding urbanization that may influence runoff, hydrologic connectivity, water chemistry, and biological conditions across both lentic and lotic habitats (Dugan et al. 2017b; Sullivan et al. 2021). We additionally calculated human-altered land use as the percentage of each 500 m buffer classified as developed or agricultural land cover using CONUS NLCD land-cover rasters. Developed land included NLCD classes 21–24, and agricultural land included classes 81–82. Collection dates were standardized to four-digit calendar years and matched to year-specific land-cover datasets. This metric was used to capture human-managed land cover that may affect aquatic habitats through diffuse inputs, altered runoff pathways, vegetation removal, soil disturbance, and landscape modification not fully represented by impervious surface alone.

### Site classification

Based on observations of flow state and system size from field sampling and Google Earth imagery, sampling sites were assigned to two functional hydrologic categories: free flowing and bounded. These categories capture major differences in hydrologic connectivity and physical constraint that could influence dispersal, disturbance, residence time, and local environmental filtering (Townsend and Hildrew 1994; Leibowitz et al. 2018, 2023). The nine sites classified as free flowing were shallow, visibly moving channels including streams, tributaries, and flow-connected wetlands with continuous or seasonally persistent surface flow and no permanent physical barrier restricting water movement at the sampling reach (Leibowitz et al. 2018). The five bounded sites generally included small ponds, isolated or semi-isolated wetlands, and small impounded tributaries with little or no surface flow and where permanent physical boundaries limited hydrologic exchange (Leibowitz et al. 2018, 2023). A notable exception was a site corresponding to a lacustrine margin within McDowell Nature Preserve along Lake Wylie (Site 12), which had a much larger surface area and internal habitat heterogeneity than the other four bounded sites. Because of these pronounced differences in areal extent and habitat scale, downstream ordination analyses comparing free flowing and bounded sites were conducted with and without Site 12.

To evaluate whether our habitat classifications corresponded to distinct environmental and landscape contexts, we compared sites in a multivariate space defined by measured environmental variables and GIS-derived covariates. Because these variables were measured on different scales, including distances, fractions, percentages, and surface-water occurrence scores, each variable was standardized across sites prior to analysis. For each variable, z-scores were calculated as the site value minus the study-wide mean divided by the study-wide standard deviation. Standardized values therefore represent each site’s relative position for a given variable, with positive values indicating above-average values, negative values indicating below-average values, and values near zero indicating conditions close to the study mean. The resulting site-by-variable matrix was used to calculate pairwise environmental dissimilarity among sites and to visualize site relationships using non-metric multidimensional scaling. We then tested whether environmental profiles differed among habitat classifications using PERMANOVA, with habitat type as the grouping factor.

### eDNA Sampling and Collection

Sampling was conducted at the 14 urban aquatic sites (Fig. 1) during fall in two consecutive years (September 19 to October 7, 2023, and from October 6 to October 14, 2024). At each site, three 1 L water samples, were filtered *in situ* to collect environmental DNA (eDNA) using a Smith-Root eDNA Sampler following a standardized field filtration protocol (Smith-Root, Vancouver, WA) and practices to prevent cross-contamination (Goldberg et al. 2016). Triplicate samples were collected using the Smith-Root trident pole for triplicate sampling as independent field filtrations from the same local habitat to capture fine-scale variation in eDNA availability while maintaining consistent sampling effort across sites. Water samples were collected from the bank, at a water depth ≥30 cm to reduce sediment entrainment from the bottom and filter clogging, with the intake positioned in the active channel of streams or in the littoral area of ponds depending on site type. Water samples were filtered through 0.5 µm pore-size filters, which were immediately sealed in their self-preserving housings after filtration. Additionally, field blanks consisting of 1 L of nuclease-free water were filtered once per sampling season at the parking lot entrance to one of the sites, Ribbonwalk Nature Preserve (site 4; Fig. 1) The blank was exposed to ambient conditions for five minutes, and handled using the same filtration procedure to assess potential contamination during sample handling and transport. At each site, water temperature, pH, conductivity, dissolved oxygen, and specific conductance were measured *in situ* to characterize local abiotic conditions. All sensors were calibrated in the lab prior to fieldwork as per the instruction manuals. We used a Divolight EZ-9909SP to capture salinity (Ppm), pH, and water temperature (°C). Dissolved oxygen (mg/L) was taken with a DO 9100 dissolved oxygen meter.

### Sequencing and Species Identification

Filtered eDNA samples were submitted to Genidaqs for DNA extraction, metabarcoding library preparation, sequencing, and initial bioinformatic processing. The laboratory workflow followed the Genidaqs vertebrate metabarcoding protocol described by Keel et al. (Keel et al. 2025). DNA was extracted from filters using the QIAamp DNA Mini Kit (QIAGEN, Maryland, United States). Extracted DNA was then purified using QIAamp spin columns (QIAGEN, Maryland, United States), passed through a Zymo OneStep PCR Inhibitor Removal Kit (Zymo Research Corporation, California, United States), and processed with extraction controls in parallel. Metabarcoding targeted a short (170 bp) fragment of the vertebrate mitochondrial 12S rRNA region using a multiplex of the MiFish-U-F and U-R primer set (Miya et al. 2015) that was amplified using PCR. Each sample was amplified in triplicate. First, PCR products were pooled and diluted, then Illumina adapter sequences were added in a second PCR, and dual-indexed barcodes were added in a final PCR. Libraries were size-selected, purified, quantified, and sequenced on an Illumina MiSeq using v2 300-cycle chemistry with a PhiX sequencing control. Environmental and laboratory controls followed Djurhuus et al. (2017) and included field blanks, extraction blanks, and PCR no-template controls.

Sequencing data were processed by Genidaqs to generate exact sequence variants (ESVs), each representing a unique DNA sequence, and assign taxonomy. Sequence data were processed using the MetaWorks pipeline with a 12S vertebrate classifier under default parameters (Keel et al. 2025). The pipeline grouped ESVs into zero-radius operational taxonomic units, quantified sequence reads per ESV in each sample, and generated provisional taxonomic identifications. ESVs represented by fewer than 100 reads were removed to reduce the influence of likely sequencing artifacts, consistent with the conservative filtering approach used by Genidaqs for ecological inference from high-read-depth metabarcoding data (Keel et al. 2025).

Provisional taxonomic assignments were verified against the NCBI GenBank nucleotide database using BLASTn or against reference mitochondrial genomes from regional species (PV446219.1 *Etheostoma flabellare* & JBAHVP010000001.1 *Fundulus rathbuni*). ESVs were assigned to species when sequence identity was at least 98% with 100% query coverage to a single species. ESVs matching multiple species at the same identity and query-coverage criteria were evaluated against regional occurrence information, and sequences that could not be resolved to a single co-occurring species were assigned to the lowest taxonomic level that included all plausible matches (e.g., “Genus”, “family”, etc). ESVs with less than 95% identity to GenBank reference sequences, or without reliable matches, were removed from the analysis. Sequences assigned to common anthropogenic sources, including human, dog, cat, cow, chicken, and pig, were removed before downstream analysis. *Mallotus villosus* (capelin) was removed from the dataset because it is a marine species not expected in freshwater North Carolina systems, and its detection likely reflects contamination from bait or food sources. Species-level assignments were refined to generate a consistent dataset aligned with NCBI taxonomy by resolving ambiguous genus-level, family-level, and dual-species annotations into single representative species. For records containing paired or uncertain identifications, assignments were standardized based on regional occurrence, ecological plausibility, and consistency with NCBI taxonomy (**see Supplemental Methods**). The resulting filtered dataset was converted to site-level vertebrate detections that were summarized as presence-absence across triplicates, and then subset to fish taxa for analyses of community composition, taxonomic diversity, functional diversity, and phylogenetic diversity. As eDNA metabarcoding detections can vary among field replicates, we assessed whether replicate samples retained consistent site-level signatures and additionally tested whether taxonomic assignments were consistent across seasons and replicates (**see Supplemental Methods; Supplemental Table 1**).

### Ecological Data

Functional ecological traits including diet and habitat use were assigned to each detected fish taxon using published natural history accounts, regional fish ecology literature, and curated trait databases such as FishBase, NatureServe, and USGS resources. When local information was unavailable, assignments were made conservatively using genus-level or species-complex ecology.

Dietary niche was characterized using a non-exclusive ordinal trait framework intended to capture dominant resource use rather than assign species to single feeding guilds. Each taxon was scored from 0 to 5 across five diet axes: piscivory, invertivory, planktivory, herbivory, and detritivory. Scores were interpreted as: 0 = no evidence of use or not reported as a diet item, 1 = rare or incidental diet item, 2 = minor or supplemental component, 3 = regular but not dominant component, 4 = major diet component for many populations or life stages, and 5 = primary or defining diet component. Because taxa could receive nonzero scores across multiple diet axes, this framework retained dietary flexibility and omnivory while allowing differences in dominant resource use to contribute to functional diversity analyses. Consumption of non-fish vertebrates, including amphibians, was treated as functionally equivalent to piscivory because these prey represent large-bodied vertebrate resources captured through similar predatory strategies. Plant material and detrital organic matter were scored separately as herbivory and detritivory to distinguish consumption of living primary producers from reliance on decomposing organic material and associated basal energy pathways. For taxa identified only to genus or species complex, scores were assigned conservatively based on feeding ecology documented for plausible regional members of the group (**Supplemental Table 2**).

Water-column habitat use was characterized using four non-exclusive vertical-position categories: benthic, demersal, pelagic, and surface. These categories were selected because they are broadly applicable across streams, wetlands, ponds, and lakes and capture major differences in where fishes forage, move, and reside within the water column. Benthic taxa were defined as species primarily associated with the substrate. Demersal taxa were defined as species that occupy or forage near the bottom but are not strictly substrate-bound. Pelagic taxa were defined as species that commonly occupy open-water habitats. Surface taxa were defined as species that frequently forage or occur near the air–water interface. Habitat-use scores were assigned on a 0–5 ordinal scale, where 0 indicates no evidence of use and 5 indicates primary or defining use of that water-column position. As with dietary traits, scores were based on typical adult ecology from published literature and databases, with conservative genus-level assignments used when species-level identifications were unavailable.

### Site-level diversity and relative-loss fingerprints

We calculated site-level biodiversity metrics across taxonomic, phylogenetic, functional, and compositional dimensions using the site-by-species presence–absence matrix, the pruned phylogeny, and the species-level trait table. Taxonomic metrics included species richness, Shannon diversity, Simpson diversity, and mean Sørensen dissimilarity to all other sites. Phylogenetic metrics used a rooted time-calibrated actinopterygian phylogeny from Rabosky et al. (2018), pruned to detected taxa represented in the dataset and included Faith’s phylogenetic diversity, mean pairwise phylogenetic distance, and mean nearest taxon distance. Functional metrics included functional richness, trait onion peeling area, functional evenness, functional divergence, and Rao’s quadratic entropy. Functional diversity metrics were calculated from the trait matrix containing body size, diet, and habitat-position traits. Diversity indices were calculated using the picante (Kembel et al. 2010), FD (Laliberté and Legendre 2010; Laliberté et al. 2014), and vegan (Oksanen et al. 2024) packages in R.

Relative-loss scores were used to construct site-level biodiversity-loss fingerprints to summarize the extent to which each site showed loss or retention across taxonomic, phylogenetic, functional, and compositional metrics. To compare biodiversity loss across metrics with different scales and units, each site-level diversity value was converted to a relative-loss score:

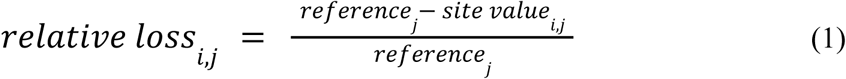

where *i* indicates site and *j* indicates diversity index. Relative-loss values were bounded from 0 to 1, with 0 indicating no loss relative to the reference value and 1 indicating complete relative loss. In this sense, “loss” refers to loss relative to the reference value used for each index, not documented historical loss as no baseline data exists for this region. Reference values were defined separately for each metric. Richness used total observed species richness across the full dataset as the reference. Shannon diversity used the log of total observed richness. Simpson diversity used 1−1/*S*, where *S* was total observed richness. Faith’s phylogenetic diversity used the total branch length of the pruned tree. Mean Sørensen dissimilarity, mean pairwise phylogenetic distance, mean nearest taxon distance, functional richness, Rao’s quadratic entropy, and trait onion peeling area used the maximum observed site value as the reference. Functional evenness and functional divergence used their theoretical maximum of 1 as the reference. All relative-loss values were visualized as a site-by-index heat map to facilitate comparison of biodiversity-loss profiles among sites.

### Ordination and clustering of site-level structure

To evaluate whether free-flowing and bounded habitats occupy distinct regions of environmental, taxonomic-composition, and biodiversity-profile space, we performed a series of ordination analyses that enable visualization of relationships among sites in abiotic variables, fish species presence and absence, and relative biodiversity-loss fingerprints. Environmental dissimilarities were calculated from a multivariate dataset of field-measured water quality variables (e.g. dissolved oxygen, salinity, and pH) and GIS-derived metrics representing hydrologic connectivity, landscape configuration, and anthropogenic land use. Because environmental variables were measured on different scales, they were standardized to z-scores (calculated as the site value minus the study-wide mean divided by the study-wide standard deviation) prior to calculating Euclidean distances among sites. Pairwise differences in fish assemblage composition were quantified using binary Jaccard dissimilarity, which measures the degree to which fish species presence-absence differs between two sites. Finally, differences among sites in multivariate biodiversity-loss profiles (represented by relative losses in taxonomic, phylogenetic, and functional diversity metrics) were calculated using the Bray-Curtis dissimilarity index.

Relationships among sampling sites in each dissimilarity matrix were visualized using non-metric multidimensional scaling (NMDS) implemented with the metaMDS function in the R package vegan (Oksanen et al. 2024). To assess the degree to which habitat type (free flowing or bounded) explained differences among sites in each ordination space, we performed permutational multivariate analysis of variance (PERMANOVA) with the adonis2 function in vegan for each dissimilarity matrix, with habitat type as the grouping variable and 999 permutations. Because PERMANOVA can confound differences in within-group dispersion with significant differences in between-group mean values, homogeneity of multivariate dispersion was evaluated for each analysis using the betadisper function, followed by permutation testing using 999 permutations.

To aid interpretation of ordination structure, variables were fit to ordination configurations using the envfit function in vegan with 999 permutations. Environmental ordinations were interpreted using fitted environmental vectors, diversity-loss ordinations using fitted diversity-index vectors, and species-composition ordinations using fitted taxon vectors. Only vectors with p<0.05 were retained for visualization. For environmental ordinations, standardized mean differences between bounded and free-flowing sites were additionally calculated for variables retained from the environmental NMDS analysis. These contrasts allowed us to inspect which habitat categories differed across a suite of hydrologic connectivity, urbanization or fragmentation, and water chemistry variables, providing support for the interpretation that bounded small and free-flowing sites represent distinct environmental contexts rather than arbitrary habitat labels. Given the contribution of urbanization-associated variables to environmental separation among sites, we also tested whether individual biodiversity indices tracked a composite urbanization axis derived from measured and GIS-based environmental variables (**See supplemental materials**). Variables used for habitat contrasts were selected from the ordination analysis, so these effect sizes were used descriptively to summarize direction and magnitude of habitat contrasts rather than as independent tests of significance.

### Partitioning of ecological and life-history traits among habitat types

To test whether habitat-associated differences were accompanied by differences in the ecological strategies represented within each local fish assemblage, we analyzed two complementary trait spaces. First, we evaluated ordinal ecological trait composition using diet and habitat-position traits, including piscivory, invertivory, planktivory, herbivory, detritivory, benthic, demersal, pelagic, and surface-use scores. Dissimilarities among sites were calculated using Gower distance with traits treated as ordered variables. Site relationships were visualized using principal coordinates analysis. Differences in trait composition among habitat categories were tested using PERMANOVA with 999 permutations, and homogeneity of multivariate dispersion was evaluated using vegan::betadisper followed by permutation testing with 999 permutations (Oksanen et al. 2024).

Second, we evaluated life-history composition. Because fecundity and reproductive age are often associated with body size (Ahti et al. 2020), we calculated log size-corrected life-history traits before aggregating these variables to the site level using phylogenetic generalized least squares with a Brownian correlation structure using the ape package in R (Paradis and Schliep 2019). Residuals from these models were retained as size-corrected fecundity and size-corrected reproductive age, respectively, and were then averaged across detected taxa within each site. To place these variables on comparable ordered scales, each variable was converted to ordered quantile bins before calculating Gower distances among sites. Principal coordinates analysis was used to visualize site relationships, and habitat-type differences were tested using the same PERMANOVA and dispersion-testing framework described above. To interpret the direction of life-history differences between habitat types, we compared site-level mean body size, mean size-corrected fecundity, and mean size-corrected reproductive age between bounded small and free-flowing sites using one-way ANOVA.

We additionally implemented a permutation-based sensitivity test to evaluate whether habitat-associated trait separation exceeded expectations given the observed incidence structure, as site-level trait composition is derived from species occurrence. Species-trait labels were randomly shuffled among taxa while holding the site-by-species incidence matrix fixed. For each permutation, site-level trait composition was recalculated and the PERMANOVA F-statistic for habitat type was recomputed. Empirical p-values were calculated as the proportion of null permutations in which *F*_null_ ≥ *F*_observed_ using 499 permutations, allowing us to test whether habitat-associated trait differences were stronger than expected from the site occupancy structure alone.

## Results & Discussion

### Free-flowing and bounded habitats represent distinct urban disturbance contexts

Environmental conditions varied substantially among sites (**Figure 1A**), with free flowing sites generally having lower oxygen levels, higher salinity, and slightly lower pH (**Figure 1B**). Together with GIS-derived landscape metrics (**Figure 1C**), these variables produced distinct environmental profiles for each site that further depicted substantial differences in abiotic characteristics between bounded and free flowing habitat types (**Figure 1D**). As the only large bounded site, McDowell Nature Preserve (Site 12) had the most unique environmental profile that dramatically differentiated it from other sites (**Supplemental Figure S2**). However, even when this site is excluded, site-level profiles nevertheless contained a strong signal of separation between free-flowing and bounded aquatic habitats (**Figure 1E**). Our NMDS analysis demonstrated bounded and free-flowing sites to occupy partially distinct regions of ordination space, with habitat type explaining significant environmental dissimilarity among sites (PERMANOVA p = 0.016 | betadisper p = 0.460). Comparisons of individual parameters and the effect-size summary revealed that this separation reflected significantly higher proportions of human-altered land use (Wilcoxon p = 0.037), impervious land cover (Wilcoxon p = 0.025), and landscape fragmentation (i.e. corridor development; Wilcoxon p = 0.025) surrounding free-flowing habitats (**Supplemental Figures S3 & S4**). On the other hand, bounded habitats exhibit significantly higher aquatic habitat availability (open water in a 1km buffer; Wilcoxon p = 0.0066) and surface water persistence (mean surface water occurrence; Wilcoxon p = 0.01), as well as lower incidence of ephemeral surface water (Wilcoxon p = 0.012; **Supplemental Figures S3 & S4**). Together, these results suggest that free-flowing and bounded habitats represent distinct environmental contexts with the potential to shape disparate faunal assemblages at fine scales within the broader Charlotte urban aquatic network.

Free-flowing systems are often treated as pathways that facilitate dispersal, recolonization, and gene flow (Tonkin et al. 2018; Prunier et al. 2023)(Campbell Grant et al. 2007; Prunier et al. 2023)(Tonkin et al. 2018; Prunier et al. 2023). This connectivity may increase the potential for recolonization of free-flowing habitats from the regional species pool, enabling these aquatic habitats to retain the biodiversity observed at a regional scale even in the face of urbanization-associated changes in abiotic conditions (Smith et al. 2015; Bourassa et al. 2017). However, this connectivity may also facilitate transmission of disturbance through urban aquatic networks, where impervious surfaces and stormwater infrastructure alter the timing, volume, and routing of runoff, changing flood frequency, sediment transport, channel stability, and habitat structure (Vietz et al. 2016; Anim and Banahene 2021; MacKenzie et al. 2022; Safdar et al. 2024). For example, urbanization has been shown to increase the frequency of bed-moving flows that alter sediment dynamics in ways that simplify channel structure (MacKenzie et al. 2022), while stormwater drainage can further amplify stream contaminant loads (Walsh et al. 2022). Indeed, in this study, the higher incidence of impervious land cover and agriculture-associated land use in the vicinity of free-flowing habitats (**Figure 1C**; **Supplemental Figs. S3 & S4**) may help to explain the significantly elevated levels of salinity observed in these habitats (**Figure 1B**; **Supplemental Figs. S3 & S4**). Taken together, our results are consistent with several hallmarks of urban stream syndrome, a recurring suite of hydrologic, geomorphic, chemical, and biological changes that emerge when urban development alters how water moves through stream networks (Walsh et al. 2005; Vietz et al. 2016; Anim and Banahene 2021). Future work should quantify the degree to which individual free-flowing sites transmit disturbance by pairing longitudinal species monitoring with storm-event hydrology and repeated water-chemistry measurements. Field assessments of bank erosion, channel incision, substrate composition, and instream habitat complexity would further help determine whether biological differences are linked to physical channel modification.

In contrast to free-flowing systems, bounded aquatic habitats are often treated as isolated or secondary water bodies within urban landscapes (Hassall 2014; Hill et al. 2017; Oertli and Parris 2019; Vasco et al. 2024). However, bounded systems are not simply disconnected versions of streams. Their physical enclosure changes the ecological consequences of urbanization by increasing the importance of water residence time, local catchment inputs, vegetation structure, management history, and episodic exchange with surrounding drainage networks (Sønderup et al. 2016; Janke et al. 2022; Marques and Mandrak 2024). For example, the degree to which stormwater ponds support freshwater biodiversity varies strongly with pond design, pollutant exposure, vegetation, hydroperiod, and landscape context (Ferzoco and McCauley 2024; McKercher et al. 2024). Similar findings have been reported for pondscapes, where local conditions can either support organismal persistence or amplify chronic filtering (Hill et al. 2021; Hamer et al. 2024; Márton et al. 2025). In short, these findings suggest that bounded habitats can retain, attenuate, or concentrate urban stressors depending on their design, hydrology, and management. Nutrients, salts, heat, contaminants, and organic matter may accumulate in poorly flushed basins (Janke et al. 2022; Izma et al. 2025), while vegetation, shallow margins, and longer residence times may create opportunities for taxa that are excluded from more frequently disturbed channels (McKercher et al. 2024; Márton et al. 2025). In our study, the environmental separation of bounded small sites from free-flowing sites suggests that these habitats are not merely endpoints along the same stream-disturbance gradient, but represent a different mode of urban aquatic habitat formation. More broadly, these results suggest that free-flowing and bounded habitats represent different ways that urban landscapes transmit, store, and transform ecological stressors.

### eDNA detections reveal a mixed urban fauna structured by habitat type

Our eDNA detections reveal that Charlotte’s urban aquatic network contains a highly mixed fauna that appears to be sorted according to local habitat context (**Figure 3**). Free-flowing sites retain a broader representation of the detected regional species pool, including shads (*Dorosoma* spp.), catfishes (e.g. *Ameiurus natalis*), suckers (e.g. *Catostomus commersonii*), minnows (e.g. *Clinostomus funduloides*), mosquitofish (*Gambusia holbrooki*), darters (*Etheostoma* sp.), bass (*Micropterus salmoides*), and sunfishes (*Lepomis* spp.; **Figure 3**). Bounded sites, on the other hand, are more dominated by lentic-associated or managed faunal elements such as sunfishes, bass, and crappies (*Pomoxis* spp.), with fewer detections of stream-associated species such as minnows, darters, and suckers (**Figure 3**). Reliability of these detections was supported by high ESV read counts of retained taxa (**Supplemental Figure S5**) and higher compositional similarity among replicates from the same site than among replicates from different sites (**Supplemental Figure S6-S10**). Ordination analyses further supported habitat-associated divergence in fish assemblages, with NMDS separating free-flowing sites dominated by stream-associated taxa from bounded sites containing more lentic, managed, and introduced assemblage elements (**Figure 4C**; PERMANOVA p = 0.001; **Supplemental Fig. S11**). These compositional differences are also reflected in taxonomic, phylogenetic, and functional diversity indices (**Figure 4A; Supplemental Table 3**). Bounded sites generally exhibited higher relative-loss values than free-flowing sites in species richness, Faith’s phylogenetic diversity, functional richness, and trait onion peeling area, indicating reduced representation of the taxonomic, phylogenetic, and functional breadth observed across the broader Charlotte metropolitan region (**Figure 4A**). This is supported by the finding that habitat types separated significantly in site-by-index relative-loss space (**Figure 4B**; PERMANOVA p = 0.039; **Supplemental Figure S11**).

**Figure 3.**
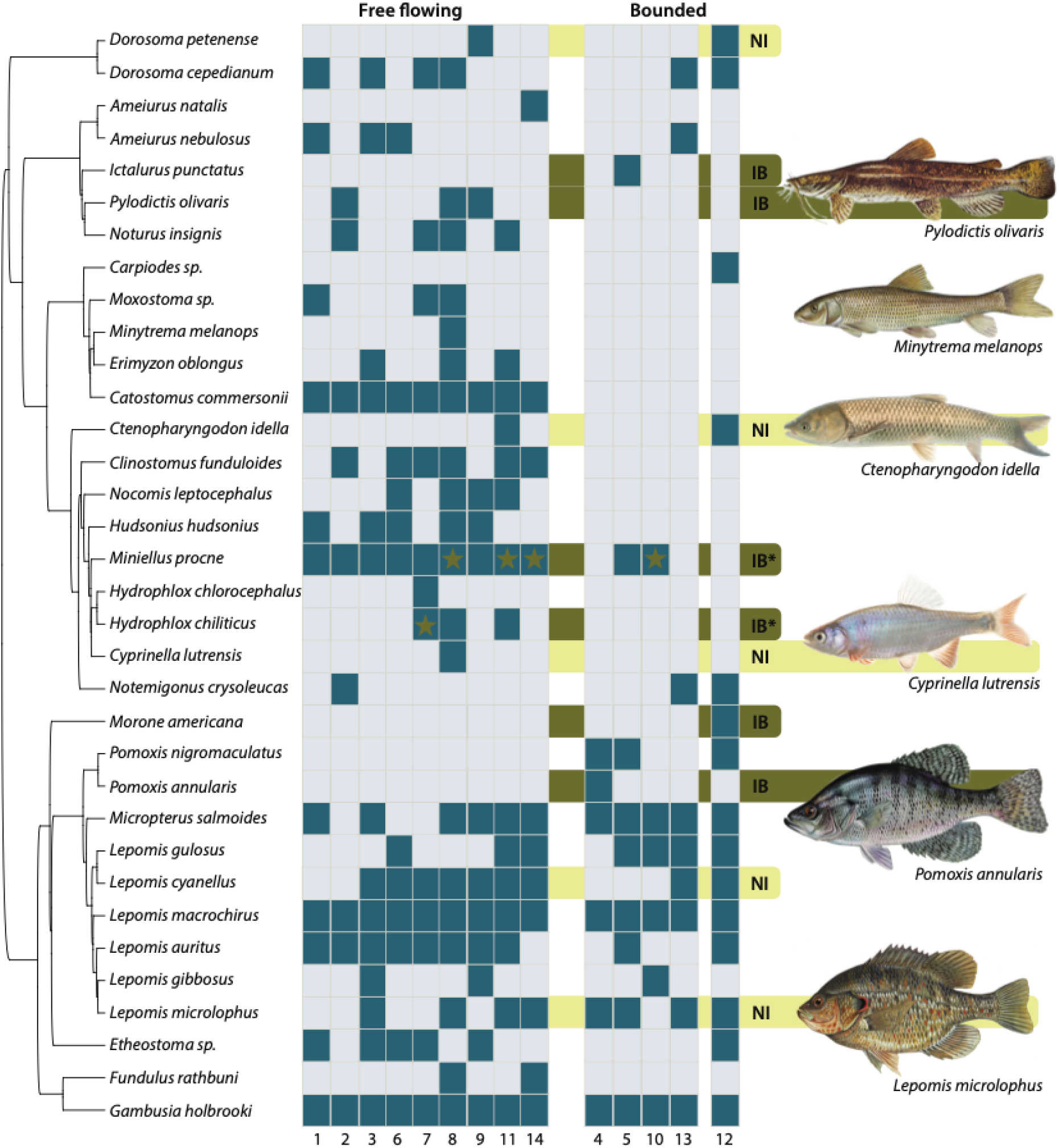
Species detected across sites. Phylogeny on the left indicates the evolutionary relationships of detected species. Cells are shaded dark for presence and light for absence with site numbers indicated in each column and corresponding to site numbers in Figure 1. Bars behind rows indicate non-indigenous (introduced; NI) or indigenous but not in this basin (IB) following Tracy et al. (Tracy et al. 2020). Stars in cells for *Miniellus procne* & *Hydrophlox chiliticus* indicate IB sites of occurrence. Representative images of fish species are from the public domain.

**Figure 4.**
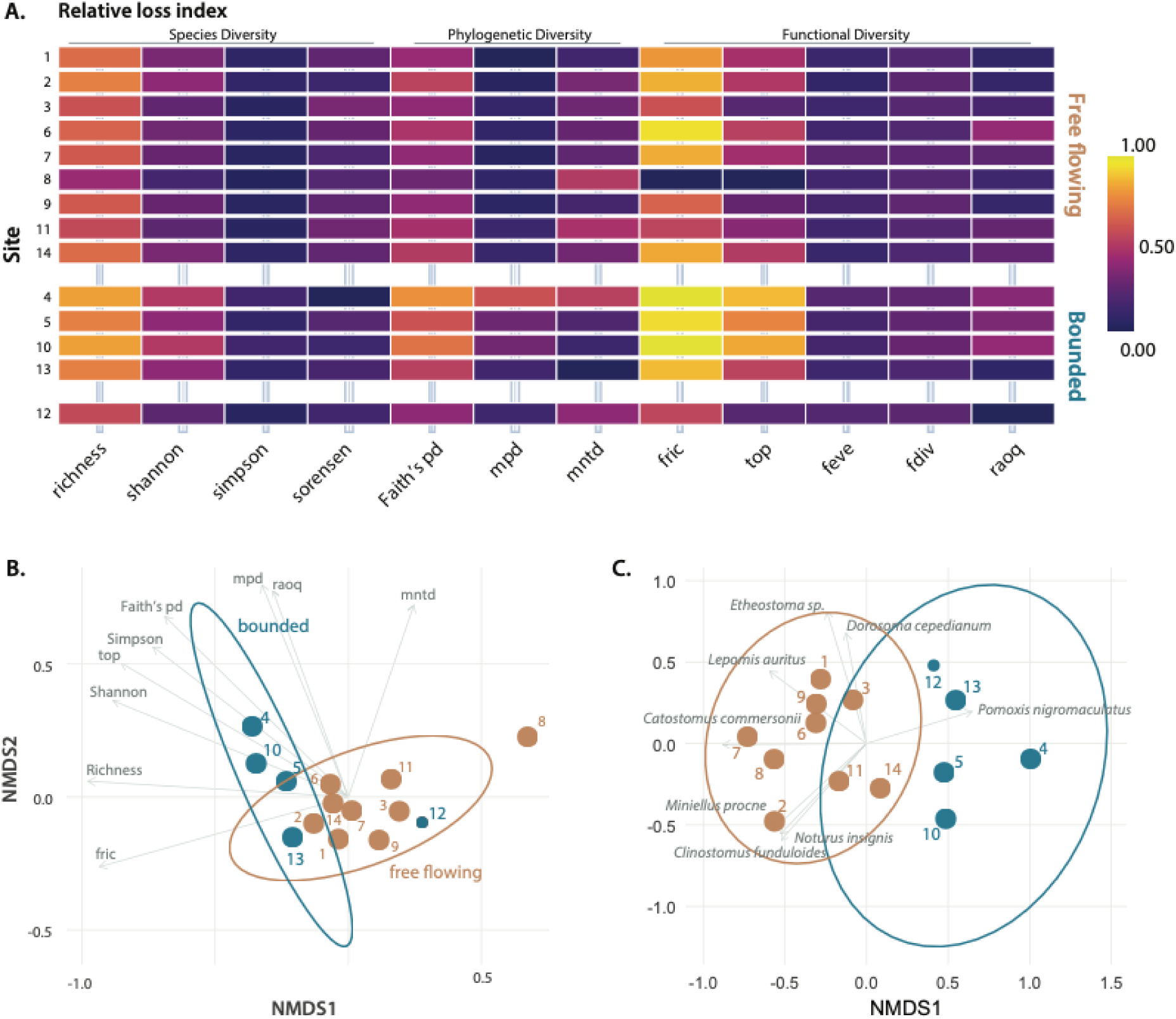
Relative diversity loss across taxonomic, phylogenetic, and functional dimensions of the Charlotte fish assemblages. **(A)** Heat map of site-level relative loss values for diversity indices, ordered by metric family and grouped by habitat type. Warmer colors indicate greater relative loss and cooler colors indicate lower relative loss. Richness = species richness, Shannon = Shannon diversity index, Simpson = Simpson diversity index, Sorensen = mean Sorensen dissimilarity to all other sites, Faith’s pd = Faith’s phylogenetic diversity, mpd = mean pairwise phylogenetic distance, mntd = mean nearest taxon phylogenetic distance, fric = functional richness, top = trait onion peeling, feve = functional evenness, fdiv = functional divergence, and raoq = Rao’s quadratic entropy. **(B)** Nonmetric multidimensional scaling (NMDS) ordination of the diversity-index loss matrix. **(C)** NMDS ordination of site-by-species presence-absence composition. Points represent sites, shadings indicate free-flowing versus bounded habitats. Ellipses summarize the free-flowing sites and the four bounded-small sites, and Site 12 remains plotted as the bounded-large comparison site.

This habitat-associated structure occurred within a fauna shaped by multiple historical, biogeographic, and management sources. More than one-third of detected taxa are either nonindigenous (NI) to North Carolina or indigenous to the state but outside their expected drainage basin (IB; (Tracy et al. 2020)). However, these detections should not be interpreted as a direct response to urbanization alone. Consistent with this interpretation, direct tests between individual biodiversity indices and the composite urbanization axis revealed limited evidence for monotonic urbanization-associated biodiversity responses across metrics (**Supplemental Figure S12; Supplemental Tables 4&5**). Instead, the species we detected are likely drawn from the native regional species pool, state-native drainage transfers, stocked or managed pond taxa, and nonindigenous generalists. This pattern is consistent with long-standing evidence that Atlantic-slope drainages in North Carolina, including the Yadkin–Pee Dee drainage basin, have accumulated nonindigenous and interbasin fish records through stocking, bait release, aquarium release, sport-fish management, and drainage-level faunal exchange (Tracy et al. 2013, 2020). Several of the taxa detected are particularly noteworthy as they represent different mechanisms by which human-mediated fish movement can alter urban aquatic assemblages. For example, Flathead Catfish (*Pylodictis olivaris*) can function as a large predatory species capable of consuming native fishes, altering predator–prey dynamics, and suppressing native fish biomass (Pine et al. 2005, 2007; Montague and Shoup 2021; Stark et al. 2024). Red-ear Sunfish (*Lepomis microlophus*) can restructure native fish abundances and resource use patterns by altering benthic food webs (Fisher Huckins et al. 2000; Whitney et al. 2021), and introductions of Grass Carp (*Ctenopharyngodon idella*) can reduce aquatic vegetation, increase turbidity, and indirectly alter habitat structure for fishes and invertebrates (Bain 1993; Dick et al. 2016; van der Lee et al. 2017). Thus introduced and interbasin taxa appear to be one component of a broader habitat-associated faunal sorting pattern rather than the sole explanation for assemblage differentiation.

Our results are consistent with recent work that highlights how urban and human-modified fish assemblages often change through nonparallel shifts in richness, species composition, functional diversity, and phylogenetic diversity. Urban streams can retain similar richness across decades even as species identities and trait structure change (Antoniazzi et al. 2023), urbanized river networks can show reduced taxonomic and functional diversity alongside increased phylogenetic diversity (Yang et al. 2023), or urban stream segments can gain species richness at the same time they lose functional richness (Qiao et al. 2022). Studies of invasion and homogenization further reinforce this point. Nonnative fishes can reduce functional diversity, reduce beta diversity, and increase assemblage similarity even when species counts alone give an incomplete picture of ecological change (Villéger et al. 2014; Toussaint et al. 2016; Moore and Olden 2017; Milardi et al. 2019; Morrill et al. 2024). The pattern we observe in Charlotte adds a habitat-scale perspective to this broader literature: urban biodiversity change in this network is better understood as habitat-specific sorting of a mixed regional species pool than as a uniform gradient of richness loss.

### Historical records define the mixed regional species pool

Placing our detections into historical context shows that the assemblages detected by eDNA embedded within a long, uneven regional sampling archive rather than standing apart from it (**Figure 5**). Historical fish records from Mecklenburg County follow the same broad temporal structure reported for North Carolina freshwater fishes as a whole, with early museum and survey collecting followed by a larger increase in records during recent decades that parallels broader expansion in environmental monitoring, agency surveys, biodiversity databasing, and conservation awareness (Boakes et al. 2010; Sigler et al. 2021; Bowler et al. 2022). These records are therefore most useful not as a standardized temporal baseline, but as a documented regional species pool against which contemporary detections can be interpreted. Our eDNA detections largely recovered abundant and repeatedly documented taxa while also detecting less frequently recorded species such as Yellow Bullhead (*Ameiurus natalis*) and several noteworthy regional records that expand the scope of occurrences from adjacent regions further into the Charlotte metropolitan area. These include multiple Flathead Catfish detections and a possible Carolina Quillback locality (**Supplemental Figures. 13-14**). Although tempting, we caution that these historical records cannot be treated as a standardized baseline for site-level temporal change because sampling locations, gears, effort, and objectives varied across decades and sites. Instead, it clarifies which taxa are consistently documented, which taxa remain sparsely represented, and how contemporary eDNA detections fit within a longer history of native persistence, drainage exchange, stocking, introduction, and incomplete sampling.

**Figure 5.**
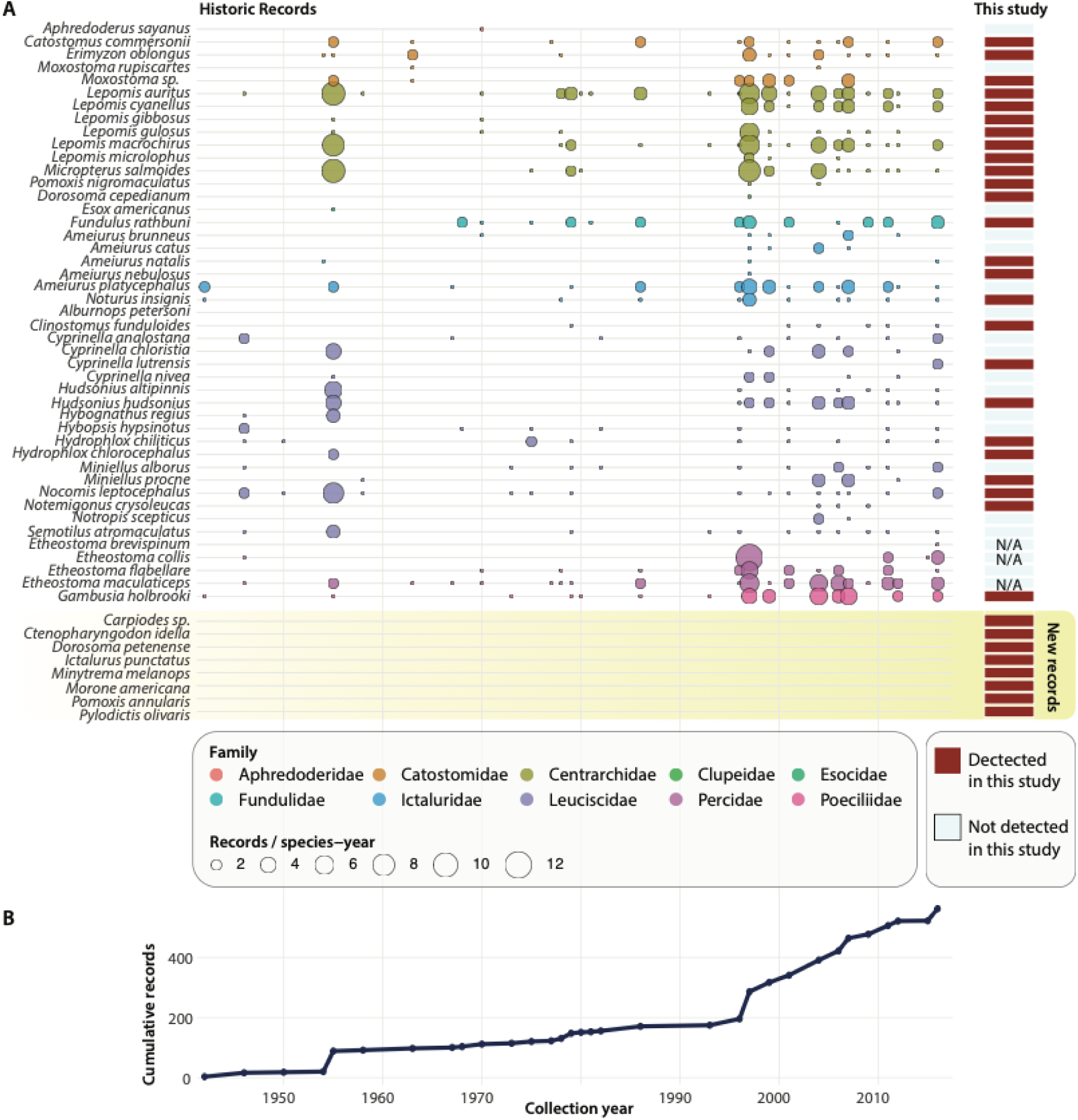
Historic fish records within the Charlotte metro study footprint and comparison with taxa detected by the current eDNA study. **(A)** Bubble timeline of historic records retained within the study convex hull plus a one-mile buffer. Bubble size represents the number of records for a species in a given collection year with shadings indicating taxonomic families. The right-side grid depicts whether each taxon was detected in the current study (dark: detected, light:not detected). The bottom shaded block highlights the seven taxa with no historic records inside of our metro study area: *Carpiodes sp.* (Carolina Quillback), *Ctenopharyngodon idella* (Grass Carp), *Dorosoma petenense* (Threadfin Shad), *Ictalurus punctatus* (Channel Catfish), *Minytrema melanops* (Spotted Sucker), *Morone americana* (White Perch), and *Pylodictis olivaris* (Flathead Catfish). **(B)** Cumulative number of historic records that have been archived within our study region through time.

This historical framing also underscores why eDNA records should be interpreted through regional faunal knowledge, taxonomic verification, and complementary field evidence. eDNA metabarcoding can extend conventional surveys by detecting fish diversity in habitats that are difficult to sample or where taxa are rare, elusive, or unevenly detected by gear (Miya et al. 2015; Gibson et al. 2023). However, metabarcoding detections depend on reference-library completeness, marker resolution, primer performance, and local species representation, which can produce ambiguous assignments or taxonomic blind spots when candidate species lack comparable genetic reference data (Evans et al. 2017; Jerde et al. 2021; Keck et al. 2023; Morey et al. 2024) This limitation was evident in our unresolved darter detections. The absence of mitochondrial reference data for several plausible candidate species prevented species-level assignment, whereas the available mitochondrial genome for Fantail Darter (*Etheostoma flabellare*) allowed that species to be ruled out as a non-match. This example echoes a growing sentiment that eDNA results are strongest as part of an integrated evidentiary framework. Museum specimens provide the material basis for resolving taxonomic uncertainty and expanding genomic databases (de Santana et al. 2021; Schmid et al. 2025), while conventional surveys provide information on abundance,verified local occurrence, demographic structure, and habitat associations. Regardless, in the context of this study, the available historical archive and eDNA detections together define the mixed regional species pool that bounded and free-flowing habitats subsequently filter into distinct assemblage-level outcomes.

### Habitat type filters assemblage-level life-history composition

Urban fish assemblages often contain species drawn from multiple biogeographic and management pathways. As a result, the exact species present at any one site may reflect historical contingency as much as contemporary habitat filtering. However, our results demonstrate that habitat type can still impose predictable structure on the kinds of life histories represented in local assemblages. In our data, ordinal ecological trait space differed significantly between habitat types without evidence of unequal dispersion, indicating a shift in the average diet and habitat-position traits represented in bounded versus free-flowing assemblages (**Figure 6A**; PERMANOVA R² = 0.350, p = 0.002; betadisper p = 0.751). Life-history space showed an even stronger habitat association, with bounded and free-flowing sites separating primarily along Axis 1 (**Figure 6B**; PERMANOVA R² = 0.632, p = 0.002; ANOSIM R = 0.797, p = 0.002; Axis 1 Wilcoxon p = 0.003). The direction of this shift was consistent across ordination vectors and site-level contrasts: bounded sites contained assemblages with larger mean body size, higher size-corrected egg production, and lower size-corrected reproductive age than free-flowing sites (**Figures 6C–E**; Wilcoxon p = 0.033, p = 0.003, and p = 0.008, respectively). The species-level occurrence models sharpen this interpretation. Habitat-by-trait interactions were not significant for log body size, egg residuals, or reproductive-age residuals after accounting for repeated observations of species and sites (interaction p = 0.517–0.993), indicating that the pattern is not a simple rule in which individual traits predict occurrence differently in each habitat type. Instead, habitat type appears to sort a historically mixed regional species pool into convergent assemblage-level life-history profiles.

**Figure 6.**
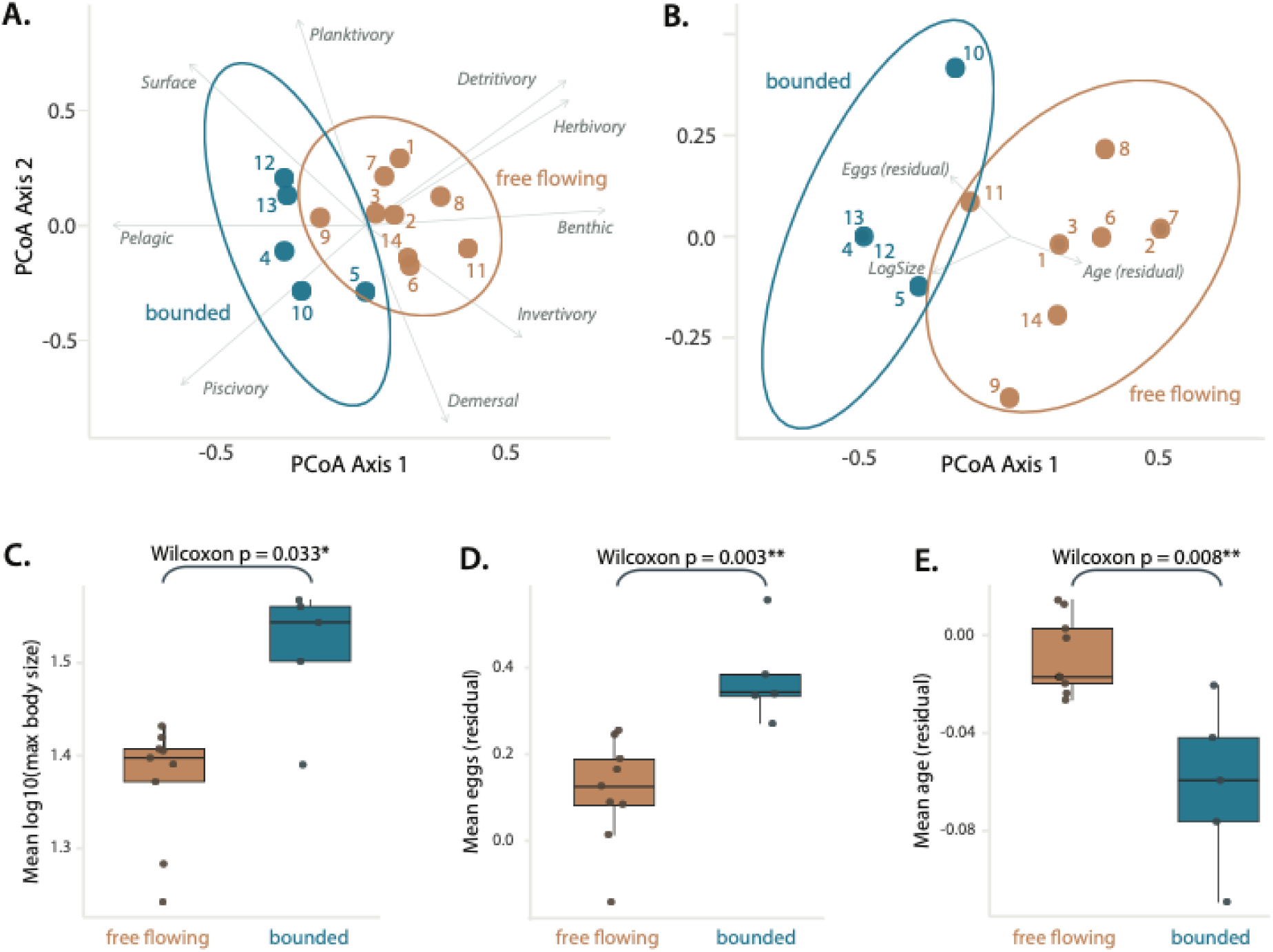
Trait structure and site-level summaries across habitat categories. (A) Principal coordinates analysis (PCoA) of ordinal trophic and water-column habitat traits based on Gower distance. (B) PCoA of categorical life-history traits, with body size and reproductive residuals binned into ordered categories. Points represent sites, colored by habitat category, with site identifiers shown. Ellipses indicate group dispersion where sample size permitted. Panel subtitles report PERMANOVA and betadisper permutation p-values. (C) Mean log10 maximum body size by habitat category. (D) Mean egg-production residual after accounting for body size. (E) Mean reproductive-age residual after accounting for body size. Boxplots show site-level values with individual sites overlaid; subtitles report Wilcoxon p-values.

Some separation between stream-associated and lentic-associated fishes is expected as flowing and standing waters often differ in current velocity, substrate, vegetation, thermal structure, predator environment, and disturbance regime (Winemiller and Rose 1992; Vila-Gispert et al. 2002; Blanck et al. 2007; Mims et al. 2010; Mims and Olden 2013). However, stream and pond faunas are not usually separated by a hard boundary. Streams contain pools, backwaters, floodplain connections, side channels, and low-velocity margins that can support taxa associated with slower water, while ponds and impoundments embedded within drainage networks can retain portions of the stream fauna, especially when exchange occurs through flooding, outlet connections, or recent formation from formerly flowing reaches (Schlosser and Kallemeyn 2000; Bower et al. 2019; O’Mara et al. 2024). This expectation is especially relevant in the southeastern United States, where beaver activity, floodplain exchange, and small impoundments create pond-like habitats within stream networks and generate substantial overlap between lotic and lentic assemblages (Snodgrass and Meffe 1998; Schlosser and Kallemeyn 2000; Kirsch and Peterson 2014). The sharp separation we observe between habitat types suggests that urban environments may be deepening the lotic–lentic life history divide.

Fish assemblages are filtered by hydrologic predictability, disturbance, and regulation, with altered or stabilized systems favoring different life-history strategies than variable, connected, or disturbance-prone channels (Poff et al. 1997; Olden et al. 2006; Perkin et al. 2017; Hitt et al. 2022). Urban systems add a further layer because imperviousness and stormwater infrastructure route rainfall rapidly into channels, increasing peak flows, stormflow volumes, sediment transport, channel erosion, and physical habitat instability in free-flowing systems (O’Driscoll et al. 2010; Violin et al. 2011) that is particularly pronounced in this region (Boggs and Sun 2011). At the same time, bounded urban waters can create slower, more persistent, and more managed aquatic environments where residence time, vegetation, stocking, predator introductions, and pond design influence which taxa establish and persist (Oertli and Parris 2019; Ferzoco and McCauley 2024; McKercher et al. 2024). If this sorting operated as a simple species-level trait rule, then body size, fecundity, or reproductive age should predict habitat occupancy differently in bounded and free-flowing sites. However, they did not. Species-level occurrence models showed no significant habitat-by-trait interactions, indicating that the strongest signal is not differential occurrence of individual species based on single life-history traits. Instead, the pattern emerges after historically contingent species pools are assembled into communities (**Figure 6**). Species composition can remain historically contingent because each site receives a different mixture of native taxa, drainage transfers, stocked species, and nonindigenous generalists, while life-history composition can still converge because bounded habitats repeatedly favor larger-bodied, more fecund, earlier-maturing profiles, while free-flowing habitats retain a different stream-associated life-history structure. More broadly, these results suggest that urban aquatic habitats can amplify selective pressures on lotic or lentic life histories, thereby homogenizing ecological strategy without homogenizing species composition.

## Conclusion

We demonstrate that the urban aquatic network of Charlotte is not organized simply by a gradient of degradation or by the presence of nonindigenous taxa alone. Free-flowing and bounded habitats represented distinct environmental contexts, with our eDNA detections revealing a fauna assembled from native drainage species, interbasin transfers, stocked or managed pond taxa, and nonindigenous generalists. These assemblages differed across taxonomic, phylogenetic, functional, and life-history dimensions, reflecting an urban aquatic system where bounded and free-flowing habitat types filter a historically mixed regional species pool into different assemblage-level outcomes. Although species identities remain contingent on drainage history, management, introductions, and local availability, bounded and free-flowing habitats nevertheless retain different ecological strategies and life-history profiles. This has significant implications for urban biodiversity monitoring as species counts and native/non-native lists can make communities appear superficially similar while missing deeper reorganization in the functions, lineages, and life histories. Overlooking these underlying shifts can lead to misinterpretations of ecological stability, mask emerging vulnerabilities, and limit the ability of conservation managers to anticipate how urban freshwater systems will respond to ongoing environmental pressures that impact how freshwater assemblages persist.

## Acknowledgements

We thank Rittika Malik, David Perrella, and Chris Avery for their help in the field collecting eDNA samples, and Jeff Jolly for his help with identifying eDNA field protocols.

## Conflict of interest

The authors report no conflict of interest.

## AI Usage Statement

Artificial intelligence tools were used in a limited capacity to assist with language editing and computational workflows. ChatGPT (OpenAI) was used for grammar, wording, and organizational suggestions during manuscript preparation. OpenAI Codex v5.4 was used to assist with code development, debugging, and code annotation. All scientific interpretation, analytical decisions, code verification, and final text revisions were performed by the authors. All code was manually reviewed and validated by the authors and is archived as documented Quarto files to support transparency and reproducibility, as described in the Data Availability Statement.

## Data Availability

All data generated during the course of this study and code used for analysis are available on Zenodo (DOI:10.5281/zenodo.22163127; https://zenodo.org/records/22163127).

## Notes

https://zenodo.org/records/22163127

### Competing Interest Statement

The authors have declared no competing interest.

## References

Ahti P.A., Kuparinen A., Uusi-Heikkilä S. 2020. Size does matter — the eco-evolutionary effects of changing body size in fish. Environ. Rev. 28:311–324.

Alberti M., Booth D., Hill K., Coburn B., Avolio C., Coe S., Spirandelli D. 2007. The impact of urban patterns on aquatic ecosystems: An empirical analysis in Puget lowland sub-basins. Landsc. Urban Plan. 80:345–361.

Albert J.S., Destouni G., Duke-Sylvester S.M., Magurran A.E., Oberdorff T., Reis R.E., Winemiller K.O., Ripple W.J. 2021. Scientists’ warning to humanity on the freshwater biodiversity crisis. Ambio. 50:85–94.

Alberts J.M., Fritz K.M., Buffam I. 2018. Response to basal resources by stream macroinvertebrates is shaped by watershed urbanization, riparian canopy cover, and season. Freshw. Sci. 37:640–652.

Anim D.O., Banahene P. 2021. Urbanization and stream ecosystems: the role of flow hydraulics towards an improved understanding in addressing urban stream degradation. Environ. Rev. 29:401–414.

Antoniazzi R., Montaña C.G., Peterson D., Schalk C.M. 2023. Changes in taxonomic and functional diversity of an urban stream fish assemblage: A 30-year perspective. Front. Environ. Sci. 10.

Bacheler N.M. 2026. Review of methods for sampling fish in structured habitats. Rev. Fish. Sci. Aquac. 34:223–257.

Bain M.B. 1993. Assessing impacts of introduced aquatic species: Grass carp in large systems. Environ. Manage. 17:211–224.

Barbarossa V., Schmitt R.J.P., Huijbregts M.A.J., Zarfl C., King H., Schipper A.M. 2020. Impacts of current and future large dams on the geographic range connectivity of freshwater fish worldwide. Proc Natl Acad Sci U S A. 117:3648–3655.

Blanck A., Tedesco P.A., Lamouroux N. 2007. Relationships between life-history strategies of European freshwater fish species and their habitat preferences. Freshw. Biol. 52:843–859.

Blowes S.A., Supp S.R., Antão L.H., Bates A., Bruelheide H., Chase J.M., Moyes F., Magurran A., McGill B., Myers-Smith I.H., Winter M., Bjorkman A.D., Bowler D.E., Byrnes J.E.K., Gonzalez A., Hines J., Isbell F., Jones H.P., Navarro L.M., Thompson P.L., Vellend M., Waldock C., Dornelas M. 2019. The geography of biodiversity change in marine and terrestrial assemblages. Science. 366:339–345.

Boakes E.H., McGowan P.J.K., Fuller R.A., Chang-qing D., Clark N.E., O’Connor K., Mace G.M. 2010. Distorted views of biodiversity: spatial and temporal bias in species occurrence data. PLoS Biol. 8:e1000385.

Boggs J.L., Sun G. 2011. Urbanization alters watershed hydrology in the Piedmont of North Carolina. Ecohydrology. 4:256–264.

Bourassa A.L., Fraser L., Beisner B.E. 2017. Benthic macroinvertebrate and fish metacommunity structure in temperate urban streams. J. Urban Ecol. 3.

Bower L.M., Keppeler F.W., Cunha E.R., Quintana Y., Saenz D.E., Lopez-Delgado E.O., Bokhutlo T., Arantes C.C., Andrade M.C., Robertson C.R., Mayes K.B., Winemiller K.O. 2019. Effects of hydrology on fish diversity and assemblage structure in a Texan coastal plains river. Trans. Am. Fish. Soc. 148:207–218.

Bowler D.E., Callaghan C.T., Bhandari N., Henle K., Benjamin Barth M., Koppitz C., Klenke R., Winter M., Jansen F., Bruelheide H., Bonn A. 2022. Temporal trends in the spatial bias of species occurrence records. Ecography (Cop.). 2022.

Brown L.R., Gregory M.B., May J.T. 2009. Relation of urbanization to stream fish assemblages and species traits in nine metropolitan areas of the United States. Urban Ecosyst. 12:391–416.

Campbell Grant E.H., Lowe W.H., Fagan W.F. 2007. Living in the branches: population dynamics and ecological processes in dendritic networks. Ecol. Lett. 10:165–175.

Cancellario T., Miranda R., Baquero E., Fontaneto D., Martínez A., Mammola S. 2022. Climate change will redefine taxonomic, functional, and phylogenetic diversity of Odonata in space and time. NPJ Biodivers. 1:1.

Centralina Regional Council. 2020. Regional Growth. Available from https://centralina.org/regional-collaboration/regional-growth/.

Charlotte U. 2026. History of Uptown. Available from https://uptowncharlotte.com/about/history-of-uptown.

Collen B., Whitton F., Dyer E.E., Baillie J.E.M., Cumberlidge N., Darwall W.R.T., Pollock C., Richman N.I., Soulsby A.-M., Böhm M. 2014. Global patterns of freshwater species diversity, threat and endemism. Glob. Ecol. Biogeogr. 23:40–51.

Copp G.H., Wesley K.J., Vilizzi L. 2005. Pathways of ornamental and aquarium fish introductions into urban ponds of Epping Forest (London, England): the human vector. J. Appl. Ichthyol. 21:263–274.

Dick G.O., Smith D.H., Schad A.N., Owens C.S. 2016. Native aquatic vegetation establishment in the presence of triploid grass carp. Lake Reserv. Manag. 32:225–233.

Dornburg A., Forrestel E., Moore J., Iglesias T., Jones A., Rao L., Warren D. 2017. An assessment of sampling biases across studies of diel activity patterns in marine ray-finned fishes (Actinopterygii). Bull. Mar. Sci. 93:611–639.

Dugan H.A., Bartlett S.L., Burke S.M., Doubek J.P., Krivak-Tetley F.E., Skaff N.K., Summers J.C., Farrell K.J., McCullough I.M., Morales-Williams A.M., Roberts D.C., Ouyang Z., Scordo F., Hanson P.C., Weathers K.C. 2017a. Salting our freshwater lakes. Proc. Natl. Acad. Sci. U. S. A. 114:4453–4458.

Dugan H.A., Summers J.C., Skaff N.K., Krivak-Tetley F.E., Doubek J.P., Burke S.M., Bartlett S.L., Arvola L., Jarjanazi H., Korponai J., Kleeberg A., Monet G., Monteith D., Moore K., Rogora M., Hanson P.C., Weathers K.C. 2017b. Long-term chloride concentrations in North American and European freshwater lakes. Sci. Data. 4:170101.

Elkins D., Sweat S.C., Kuhajda B.R., George A.L., Hill K.S., Wenger S.J. 2019. Illuminating hotspots of imperiled aquatic biodiversity in the southeastern US. Glob. Ecol. Conserv. 19:e00654.

Evans N.T., Li Y., Renshaw M.A., Olds B.P., Deiner K., Turner C.R., Jerde C.L., Lodge D.M., Lamberti G.A., Pfrender M.E. 2017. Fish community assessment with eDNA metabarcoding: effects of sampling design and bioinformatic filtering. Can. J. Fish. Aquat. Sci. 74:1362–1374.

Ferzoco I.M.C., McCauley S.J. 2024. Novel habitats for biodiversity? A systematic review and meta-analysis of freshwater biodiversity in stormwater management ponds. Sci Total Environ. 942:173467.

Fischer J.R., Quist M.C. 2014. Gear and seasonal bias associated with abundance and size structure estimates for lentic freshwater fishes. J. Fish Wildl. Manag. 5:394–412.

Fisher Huckins C.J., Osenberg C.W., Mittelbach G.G. 2000. Species introductions and their ecological consequences: an example with congeneric sunfish. Ecol. Appl. 10:612–625.

Fuller M.R., Doyle M.W., Strayer D.L. 2015. Causes and consequences of habitat fragmentation in river networks. Ann N Y Acad Sci. 1355:31–51.

Gibson T.I., Carvalho G., Ellison A., Gargiulo E., Hatton-Ellis T., Lawson-Handley L., Mariani S., Collins R.A., Sellers G., Antonio Distaso M., Zampieri C., Creer S. 2023. Environmental DNA metabarcoding for fish diversity assessment in a macrotidal estuary: A comparison with established fish survey methods. Estuar. Coast. Shelf Sci. 294:108522.

Goldberg C.S., Turner C.R., Deiner K., Klymus K.E., Thomsen P.F., Murphy M.A., Spear S.F., McKee A., Oyler-McCance S.J., Cornman R.S., Laramie M.B., Mahon A.R., Lance R.F., Pilliod D.S., Strickler K.M., Waits L.P., Fremier A.K., Takahara T., Herder J.E., Taberlet P. 2016. Critical considerations for the application of environmental DNA methods to detect aquatic species. Methods Ecol. Evol. 7:1299–1307.

Haddad N.M., Brudvig L.A., Clobert J., Davies K.F., Gonzalez A., Holt R.D., Lovejoy T.E., Sexton J.O., Austin M.P., Collins C.D., Cook W.M., Damschen E.I., Ewers R.M., Foster B.L., Jenkins C.N., King A.J., Laurance W.F., Levey D.J., Margules C.R., Melbourne B.A., Nicholls A.O., Orrock J.L., Song D.-X., Townshend J.R. 2015. Habitat fragmentation and its lasting impact on Earth’s ecosystems. Sci Adv. 1:e1500052.

Hamer A.J., Barta B., Márton Z., Vad C.F., Szabó B., Tornero I., Horváth Z. 2024. Patterns and correlates in the distribution, design and management of garden ponds along an urban–rural gradient. Urban Ecosyst. 27:1915–1930.

Hassall C. 2014. The ecology and biodiversity of urban ponds. WIREs Water. 1:187–206.

Hilburn B.G., Rider S.J., Johnston C.E. 2025. Biogeographic considerations for fish-based indices of stream health in regions with high species richness and endemism: A perspective from the southeastern US. Environ. Manage. 75:167–174.

Hillebrand H., Blasius B., Borer E.T., Chase J.M., Downing J.A., Eriksson B.K., Filstrup C.T., Harpole W.S., Hodapp D., Larsen S., Lewandowska A.M., Seabloom E.W., Van de Waal D.B., Ryabov A.B. 2018. Biodiversity change is uncoupled from species richness trends: Consequences for conservation and monitoring. J. Appl. Ecol. 55:169–184.

Hill M.J., Biggs J., Thornhill I., Briers R.A., Gledhill D.G., White J.C., Wood P.J., Hassall C. 2017. Urban ponds as an aquatic biodiversity resource in modified landscapes. Glob. Chang. Biol. 23:986–999.

Hill M.J., Wood P.J., Fairchild W., Williams P., Nicolet P., Biggs J. 2021. Garden pond diversity: Opportunities for urban freshwater conservation. Basic Appl. Ecol. 57:28–40.

Hitt N.P., Landsman A.P., Raesly R.L. 2022. Life history strategies of stream fishes linked to predictors of hydrologic stability. Ecol Evol. 12:e8861.

Hughes K. 2021. The world’s forgotten fishes. .

Izma G., Raby M., Ijzerman M., Prosser R., Helm P., Renaud J., Sumarah M., McIsaac D., Rooney R. 2025. An urban stormwater contaminant signature: Defining priority contaminants for urban stormwater research. Water Environ. Res. 97:e70150.

Janke B.D., Finlay J.C., Taguchi V.J., Gulliver J.S. 2022. Hydrologic processes regulate nutrient retention in stormwater detention ponds. Sci. Total Environ. 823:153722.

Jerde C.L., Mahon A.R., Campbell T., McElroy M.E., Pin K., Childress J.N., Armstrong M.N., Zehnpfennig J.R., Kelson S.J., Koning A.A., Ngor P.B., Nuon V., So N., Chandra S., Hogan Z.S. 2021. Are genetic reference libraries sufficient for environmental DNA metabarcoding of Mekong River Basin fish? Water (Basel). 13:1767.

Karr J.R. 1981. Assessment of Biotic Integrity Using Fish Communities. Fisheries. 6:21–27.

Keck F., Couton M., Altermatt F. 2023. Navigating the seven challenges of taxonomic reference databases in metabarcoding analyses. Mol Ecol Resour. 23:742–755.

Keel D.J., Karpenko K., Blankenship S.M., Schumer G., O’Rourke O., Ostberg C.O., Chase D.A., Duda J.J. 2025. A molecular specimen bank for contemporary and future study captures landscape-scale biodiversity baselines before Klamath River dam removal. Sci Rep. 15:20679.

Kembel S.W., Cowan P.D., Helmus M.R., Cornwell W.K., Morlon H., Ackerly D.D., Blomberg S.P., Webb C.O. 2010. Picante: R tools for integrating phylogenies and ecology. Bioinformatics. 26:1463–1464.

Kirsch J.E., Peterson J.T. 2014. A multi-scaled approach to evaluating the fish assemblage structure within southern Appalachian streams. Trans. Am. Fish. Soc. 143:1358–1371.

Kondratyeva A., Knapp S., Durka W., Kühn I., Vallet J., Machon N., Martin G., Motard E., Grandcolas P., Pavoine S. 2020. Urbanization effects on biodiversity revealed by a two-scale analysis of species functional uniqueness vs. Redundancy. Front. Ecol. Evol. 8.

Kraft N.J.B., Comita L.S., Chase J.M., Sanders N.J., Swenson N.G., Crist T.O., Stegen J.C., Vellend M., Boyle B., Anderson M.J., Cornell H.V., Davies K.F., Freestone A.L., Inouye B.D., Harrison S.P., Myers J.A. 2011. Disentangling the drivers of β diversity along latitudinal and elevational gradients. Science. 333:1755–1758.

Kwik J., Kho Z.Y., Quek B.S., Tan H.H., Yeo D. 2013. Urban stormwater ponds in Singapore: potential pathways for spread of alien freshwater fishes. Bioinvasions Rec. 2:239–245.

Laliberté E., Legendre P. 2010. A distance-based framework for measuring functional diversity from multiple traits. Ecology. 91:299–305.

Laliberté E., Legendre P., Shipley B. 2014. FD: measuring functional diversity from multiple traits, and other tools for functional ecology. .

van der Lee A.S., Johnson T.B., Koops M.A. 2017. Bioenergetics modelling of grass carp: Estimated individual consumption and population impacts in Great Lakes wetlands. J. Great Lakes Res. 43:308–318.

LeGrand, H., J. Amoroso, and T. Howard. 2026. Freshwater fishes of North Carolina. Available from https://auth1.dpr.ncparks.gov/fish/index.php.

Lehner B., Grill G. 2013. Global river hydrography and network routing: baseline data and new approaches to study the world’s large river systems. Hydrol. Process. 27:2171–2186.

Leibold M.A., Holyoak M., Mouquet N., Amarasekare P., Chase J.M., Hoopes M.F., Holt R.D., Shurin J.B., Law R., Tilman D., Loreau M., Gonzalez A. 2004. The metacommunity concept: a framework for multi-scale community ecology. Ecol. Lett. 7:601–613.

Leibowitz S.G., Hill R.A., Creed I.F., Compton J.E., Golden H.E., Weber M.H., Rains M.C., Jones C.E. Jr, Lee E.H., Christensen J.R., Bellmore R.A., Lane C.R. 2023. National hydrologic connectivity classification links wetlands with stream water quality. Nat Water. 1:370–380.

Leibowitz S.G., Wigington P.J. Jr, Schofield K.A., Alexander L.C., Vanderhoof M.K., Golden H.E. 2018. CONNECTIVITY OF STREAMS AND WETLANDS TO DOWNSTREAM WATERS: AN INTEGRATED SYSTEMS FRAMEWORK. J Am Water Resour Assoc. 54:298–322.

Leprieur F., Beauchard O., Blanchet S., Oberdorff T., Brosse S. 2008. Fish invasions in the world’s river systems: when natural processes are blurred by human activities. PLoS Biol. 6:e28.

Li D., Olden J.D., Lockwood J.L., Record S., McKinney M.L., Baiser B. 2020. Changes in taxonomic and phylogenetic diversity in the Anthropocene. Proc Biol Sci. 287:20200777.

Love S.A., Lederman N.J., Widloe T., Whitledge G.W. 2019. Sources of Bighead Carp and Silver Carp found in Chicago Urban Fishing Program ponds. Trans. Am. Fish. Soc. 148:417–425.

MacKenzie K.M., Singh K., Binns A.D., Whiteley H.R., Gharabaghi B. 2022. Effects of urbanization on stream flow, sediment, and phosphorous regime. J. Hydrol. (Amst.). 612:128283.

Magurran A.E., Deacon A.E., Moyes F., Shimadzu H., Dornelas M., Phillip D.A.T., Ramnarine I.W. 2018. Divergent biodiversity change within ecosystems. Proc Natl Acad Sci U S A. 115:1843–1847.

Marques P., Mandrak N.E. 2024. Ecosystem functions in urban stormwater management ponds: A scoping review. Sustainability. 16:7766.

Márton Z., Barta B., Vad C.F., Szabó B., Hamer A.J., Kardos V., Laskai C., Fierpasz Á., Horváth Z. 2025. Effects of urbanisation, habitat characteristics, and management on garden pond biodiversity: Findings from a large-scale citizen science survey. Landsc. Urban Plan. 257:105299.

McElroy M.E., Dressler T.L., Titcomb G.C., Wilson E.A., Deiner K., Dudley T.L., Eliason E.J., Evans N.T., Gaines S.D., Lafferty K.D., Lamberti G.A., Li Y., Lodge D.M., Love M.S., Mahon A.R., Pfrender M.E., Renshaw M.A., Selkoe K.A., Jerde C.L. 2020. Calibrating environmental DNA metabarcoding to conventional surveys for measuring fish species richness. Front. Ecol. Evol. 8.

McKercher L.J., Kimball M.E., Scaroni A.E., White S.A., Strosnider W.H.J. 2024. Stormwater ponds serve as variable quality habitat for diverse taxa. Wetl. Ecol. Manag. 32:109–131.

Meador M.R. 2020. Historical changes in fish communities in urban streams of the south-eastern United States and the relative importance of water-quality stressors. Ecol. Freshw. Fish. 29:156–169.

Mehdi H., Lau S.C., Synyshyn C., Salena M.G., Morphet M.E., Hamilton J., Muzzatti M.N., McCallum E.S., Midwood J.D., Balshine S. 2021. A comparison of passive and active gear in fish community assessments in summer versus winter. Fish. Res. 242:106016.

Milardi M., Gavioli A., Soininen J., Castaldelli G. 2019. Exotic species invasions undermine regional functional diversity of freshwater fish. Sci Rep. 9:17921.

Mims M.C., Olden J.D. 2013. Fish assemblages respond to altered flow regimes via ecological filtering of life history strategies. Freshw. Biol. 58:50–62.

Mims M.C., Olden J.D., Shattuck Z.R., Poff N.L. 2010. Life history trait diversity of native freshwater fishes in North America. Ecol. Freshw. Fish. 19:390–400.

Miranda R., Miqueleiz I. 2021. Ecology and conservation of freshwater fishes biodiversity: We need more knowledge to develop conservation strategies. Water (Basel). 13:1929.

Mittelbach G.G., Schemske D.W. 2015. Ecological and evolutionary perspectives on community assembly. Trends Ecol Evol. 30:241–247.

Miya M., Sato Y., Fukunaga T., Sado T., Poulsen J.Y., Sato K., Minamoto T., Yamamoto S., Yamanaka H., Araki H., Kondoh M., Iwasaki W. 2015. MiFish, a set of universal PCR primers for metabarcoding environmental DNA from fishes: detection of more than 230 subtropical marine species. R Soc Open Sci. 2:150088.

Montague G.F., Shoup D.E. 2021. Two decades of advancement in Flathead Catfish research. N. Am. J. Fish. Manag. 41.

Moore J.W., Olden J.D. 2017. Response diversity, nonnative species, and disassembly rules buffer freshwater ecosystem processes from anthropogenic change. Glob Chang Biol. 23:1871–1880.

Morey K.C., Myler E., Hanner R., Tetreault G. 2024. Taxonomic blind spots: A limitation of environmental DNA metabarcoding-based detection for Canadian freshwater fishes. Environ. DNA. 6.

Mori A.S., Isbell F., Seidl R. 2018. β-Diversity, Community Assembly, and Ecosystem Functioning. Trends Ecol Evol. 33:549–564.

Morrill D.P., Main J., Davenport G., Adams G.L., Adams S.R. 2024. Taxonomic and functional homogenization of fish assemblages in an Ozark river associated with pasture land use and constructed water bodies. Ecol. Freshw. Fish.

Myers J.A., Chase J.M., Jiménez I., Jørgensen P.M., Araujo-Murakami A., Paniagua-Zambrana N., Seidel R. 2013. Beta-diversity in temperate and tropical forests reflects dissimilar mechanisms of community assembly. Ecol Lett. 16:151–157.

NC Department of Environmental Quality. Public Wetlands Map. Available from https://ncdenr.maps.arcgis.com/apps/webappviewer/index.html?id=7641d40a498c47f9aa1153cdd5f63969.

Nedd R., Anandhi A. 2022. Land use changes in the southeastern United States: Quantitative changes, drivers, and expected environmental impacts. Land (Basel). 11:2246.

Newbold T., Hudson L.N., Hill S.L.L., Contu S., Lysenko I., Senior R.A., Börger L., Bennett D.J., Choimes A., Collen B., Day J., De Palma A., Díaz S., Echeverria-Londoño S., Edgar M.J., Feldman A., Garon M., Harrison M.L.K., Alhusseini T., Ingram D.J., Itescu Y., Kattge J., Kemp V., Kirkpatrick L., Kleyer M., Correia D.L.P., Martin C.D., Meiri S., Novosolov M., Pan Y., Phillips H.R.P., Purves D.W., Robinson A., Simpson J., Tuck S.L., Weiher E., White H.J., Ewers R.M., Mace G.M., Scharlemann J.P.W., Purvis A. 2015. Global effects of land use on local terrestrial biodiversity. Nature. 520:45–50.

O’Driscoll M., Clinton S., Jefferson A., Manda A., McMillan S. 2010. Urbanization effects on watershed hydrology and in-stream processes in the southern United States. Water (Basel). 2:605–648.

Oertli B., Parris K.M. 2019. Review: Toward management of urban ponds for freshwater biodiversity. Ecosphere. 10.

Oksanen J., Simpson G.L., Blanchet F.G., Kindt R., Legendre P., Minchin P.R., O’Hara R.B., Solymos P., Stevens M.H.H., Szoecs E., Wagner H., Barbour M., Bedward M., Bolker B., Borcard D., Carvalho G., Chirico M., De Caceres M., Durand S., Evangelista H.B.A., FitzJohn R., Friendly M., Furneaux B., Hannigan G., Hill M.O., Lahti L., McGlinn D., Ouellette M.-H., Ribeiro Cunha E., Smith T., Stier A., Ter Braak C.J.F., Weedon J. 2024. vegan: Community Ecology Package. .

Olden J.D., Poff N.L., Bestgen K.R. 2006. Life-history strategies predict fish invasions and extirpations in the Colorado river basin. Ecol. Monogr. 76:25–40.

O’Mara K., Venarsky M., Stewart-Koster B., McGregor G.B., Schulz C., Marshall J., Bunn S.E. 2024. Hydrological connectivity and environment characteristics explain spatial variation in fish assemblages in a wet–dry tropical river. Hydrobiologia. 851:5207–5221.

Paradis E., Schliep K. 2019. ape 5.0: an environment for modern phylogenetics and evolutionary analyses in R. Bioinformatics. 35:526–528.

Pekel J.-F., Cottam A., Gorelick N., Belward A.S. 2016. High-resolution mapping of global surface water and its long-term changes. Nature. 540:418–422.

Perkin J.S., Knorp N.E., Boersig T.C., Gebhard A.E., Hix L.A., Johnson T.C. 2017. Life history theory predicts long-term fish assemblage response to stream impoundment. Can. J. Fish. Aquat. Sci. 74:228–239.

Piano E., Souffreau C., Merckx T., Baardsen L.F., Backeljau T., Bonte D., Brans K.I., Cours M., Dahirel M., Debortoli N., Decaestecker E., De Wolf K., Engelen J.M.T., Fontaneto D., Gianuca A.T., Govaert L., Hanashiro F.T.T., Higuti J., Lens L., Martens K., Matheve H., Matthysen E., Pinseel E., Sablon R., Schön I., Stoks R., Van Doninck K., Van Dyck H., Vanormelingen P., Van Wichelen J., Vyverman W., De Meester L., Hendrickx F. 2020. Urbanization drives cross-taxon declines in abundance and diversity at multiple spatial scales. Glob Chang Biol. 26:1196–1211.

Pine W.E. III, Kwak T.J., Rice J.A. 2007. Modeling management scenarios and the effects of an introduced apex predator on a coastal riverine fish community. Trans. Am. Fish. Soc. 136:105–120.

Pine W.E. III, Kwak T.J., Waters D.S., Rice J.A. 2005. Diet selectivity of introduced Flathead catfish in coastal rivers. Trans. Am. Fish. Soc. 134:901–909.

Poff N.L., Allan J.D., Bain M.B., Karr J.R., Prestegaard K.L., Richter B.D., Sparks R.E., Stromberg J.C. 1997. The natural flow regime. Bioscience. 47:769–784.

Prunier J.G., Loot G., Veyssiere C., Poulet N., Blanchet S. 2023. Novel operational index reveals rapid recovery of genetic connectivity in freshwater fish species after riverine restoration. Conserv. Lett. 16.

Qiao J., Liu Y., Fu H., Chu L., Yan Y. 2022. Urbanization affects the taxonomic and functional alpha and beta diversity of fish assemblages in streams of subtropical China. Ecol. Indic. 144:109441.

Rabosky D.L., Chang J., Title P.O., Cowman P.F., Sallan L., Friedman M., Kaschner K., Garilao C., Near T.J., Coll M., Alfaro M.E. 2018. An inverse latitudinal gradient in speciation rate for marine fishes. Nature. 559:392–395.

Reid A.J., Carlson A.K., Creed I.F., Eliason E.J., Gell P.A., Johnson P.T.J., Kidd K.A., MacCormack T.J., Olden J.D., Ormerod S.J., Smol J.P., Taylor W.W., Tockner K., Vermaire J.C., Dudgeon D., Cooke S.J. 2019. Emerging threats and persistent conservation challenges for freshwater biodiversity. Biol Rev Camb Philos Soc. 94:849–873.

Roy A.H., Rosemond A.D., Paul M.J., Leigh D.S., Wallace J.B. 2003. Stream macroinvertebrate response to catchment urbanisation (Georgia, U.S.A.). Freshw. Biol. 48:329–346.

Safdar S., Jefferson A.J., Costello D.M., Blinn A. 2024. Urbanization and suspended sediment transport dynamics: A comparative study of watersheds with varying degree of urbanization using concentration-discharge hysteresis. ACS ES T Water. 4:3904–3917.

de Santana C.D., Parenti L.R., Dillman C.B., Coddington J.A., Bastos D.A., Baldwin C.C., Zuanon J., Torrente-Vilara G., Covain R., Menezes N.A., Datovo A., Sado T., Miya M. 2021. The critical role of natural history museums in advancing eDNA for biodiversity studies: a case study with Amazonian fishes. Sci Rep. 11:18159.

Sayer C.A., Fernando E., Jimenez R.R., Macfarlane N.B.W., Rapacciuolo G., Böhm M., Brooks T.M., Contreras-MacBeath T., Cox N.A., Harrison I., Hoffmann M., Jenkins R., Smith K.G., Vié J.-C., Abbott J.C., Allen D.J., Allen G.R., Barrios V., Boudot J.-P., Carrizo S.F., Charvet P., Clausnitzer V., Congiu L., Crandall K.A., Cumberlidge N., Cuttelod A., Dalton J., Daniels A.G., De Grave S., De Knijf G., Dijkstra K.-D.B., Dow R.A., Freyhof J., García N., Gessner J., Getahun A., Gibson C., Gollock M.J., Grant M.I., Groom A.E.R., Hammer M.P., Hammerson G.A., Hilton-Taylor C., Hodgkinson L., Holland R.A., Jabado R.W., Juffe Bignoli D., Kalkman V.J., Karimov B.K., Kipping J., Kottelat M., Lalèyè P.A., Larson H.K., Lintermans M., Lozano F., Ludwig A., Lyons T.J., Máiz-Tomé L., Molur S., Ng H.H., Numa C., Palmer-Newton A.F., Pike C., Pippard H.E., Polaz C.N.M., Pollock C.M., Raghavan R., Rand P.S., Ravelomanana T., Reis R.E., Rigby C.L., Scott J.A., Skelton P.H., Sloat M.R., Snoeks J., Stiassny M.L.J., Tan H.H., Taniguchi Y., Thorstad E.B., Tognelli M.F., Torres A.G., Torres Y., Tweddle D., Watanabe K., Westrip J.R.S., Wright E.G.E., Zhang E., Darwall W.R.T. 2025. One-quarter of freshwater fauna threatened with extinction. Nature. 638:138–145.

Schiff R., Benoit G. 2007. Effects of impervious cover at multiple spatial scales on coastal watershed streams^1^. J. Am. Water Resour. Assoc. 43:712–730.

Schlosser I.J., Kallemeyn L.W. 2000. Spatial variation in fish assemblages across a beaver-influenced successional landscape. Ecology. 81:1371–1382.

Schmid S., Straube N., Albouy C., Delling B., Maclaine J., Matschiner M., Møller P.R., Nocita A., Palandačić A., Rüber L., Sonnewald M., Alvarez N., Manel S., Pellissier L. 2025. Unlocking natural history collections to improve eDNA reference databases and biodiversity monitoring. Bioscience. 75:1083–1095.

Shaw J.L.A., Clarke L.J., Wedderburn S.D., Barnes T.C., Weyrich L.S., Cooper A. 2016. Comparison of environmental DNA metabarcoding and conventional fish survey methods in a river system. Biol. Conserv. 197:131–138.

Sigler K., Warren D., Tracy B., Forrestel E., Hogue G., Dornburg A. 2021. Assessing temporal biases across aggregated historical spatial data: a case study of North Carolina’s freshwater fishes. Ecosphere. 12.

Smith R.F., Venugopal P.D., Baker M.E., Lamp W.O. 2015. Habitat filtering and adult dispersal determine the taxonomic composition of stream insects in an urbanizing landscape. Freshw. Biol. 60:1740–1754.

Snodgrass J.W., Meffe G.K. 1998. Influence of beavers on stream fish assemblages:Effects of pond age and watershed position. Ecology. 79:928–942.

Sønderup M.J., Egemose S., Hansen A.S., Grudinina A., Madsen M.H., Flindt M.R. 2016. Factors affecting retention of nutrients and organic matter in stormwater ponds. Ecohydrology. 9:796–806.

Stark S., Schall M.K., Smith G.D., Maloy A.P., Coombs J.A., Wagner T., Avery J. 2024. Feeding habits and ecological implications of the invasive Flathead Catfish in the Susquehanna River basin, Pennsylvania. Trans. Am. Fish. Soc. 153:591–610.

Stoeckle M.Y., Soboleva L., Charlop-Powers Z. 2017. Aquatic environmental DNA detects seasonal fish abundance and habitat preference in an urban estuary. PLoS One. 12:e0175186.

Sullivan S.M.P., Corra J.W., Hayes J.T. 2021. Urbanization mediates the effects of water quality and climate on a model aerial insectivorous bird. Ecol. Monogr. 91:e01442.

Terando A.J., Costanza J., Belyea C., Dunn R.R., McKerrow A., Collazo J.A. 2014. The southern megalopolis: using the past to predict the future of urban sprawl in the Southeast U.S. PLoS One. 9:e102261.

Thompson P.L., Guzman L.M., De Meester L., Horváth Z., Ptacnik R., Vanschoenwinkel B., Viana D.S., Chase J.M. 2020. A process-based metacommunity framework linking local and regional scale community ecology. Ecol Lett. 23:1314–1329.

Tickner D., Opperman J.J., Abell R., Acreman M., Arthington A.H., Bunn S.E., Cooke S.J., Dalton J., Darwall W., Edwards G., Harrison I., Hughes K., Jones T., Leclère D., Lynch A.J., Leonard P., McClain M.E., Muruven D., Olden J.D., Ormerod S.J., Robinson J., Tharme R.E., Thieme M., Tockner K., Wright M., Young L. 2020. Bending the Curve of Global Freshwater Biodiversity Loss: An Emergency Recovery Plan. Bioscience. 70:330–342.

Tonkin J.D., Altermatt F., Finn D.S., Heino J., Olden J.D., Pauls S.U., Lytle D.A. 2018. The role of dispersal in river network metacommunities: Patterns, processes, and pathways. Freshw. Biol. 63:141–163.

Toussaint A., Beauchard O., Oberdorff T., Brosse S., Villéger S. 2016. Worldwide freshwater fish homogenization is driven by a few widespread non-native species. Biol. Invasions. 18:1295–1304.

Townsend C.R., Hildrew A.G. 1994. Species traits in relation to a habitat templet for river systems. Freshw. Biol. 31:265–275.

Tracy B.H., Jenkins R.E., Starnes W.C. 2013. History of fish investigations in the Yadkin–Pee Dee river drainage of North Carolina and Virginia with an analysis of nonindigenous species and invasion dynamics of three species of suckers (catostomidae). J. North Carol. Acad. Sci. 129:82–106.

Tracy B., Rohde F., Hogue G. 2020. An annotated atlas of the Freshwater Fishes of North Carolina. Southeast. Fishes Counc. Proc. 60.

Van Metre P.C., Waite I.R., Qi S., Mahler B., Terando A., Wieczorek M., Meador M., Bradley P., Journey C., Schmidt T., Carlisle D. 2019. Projected urban growth in the southeastern USA puts small streams at risk. PLoS One. 14:e0222714.

Vasco F., Perrin J.-A., Oertli B. 2024. Urban pondscape connecting people with nature and biodiversity in a medium-sized European city (Geneva, Switzerland). Urban Ecosyst. 27:1117–1137.

Vellend M. 2010. Conceptual synthesis in community ecology. Q Rev Biol. 85:183–206.

Vietz G.J., Walsh C.J., Fletcher T.D. 2016. Urban hydrogeomorphology and the urban stream syndrome. Prog. Phys. Geogr. 40:480–492.

Vila-Gispert A., Moreno-Amich R., García-Berthou E. 2002. Gradients of life-history variation: an intercontinental comparison of fishes. Rev. Fish Biol. Fish. 12:417–427.

Villéger S., Grenouillet G., Brosse S. 2014. Functional homogenization exceeds taxonomic homogenization amongEuropean fish assemblages. Glob. Ecol. Biogeogr. 23:1450–1460.

Violin C.R., Cada P., Sudduth E.B., Hassett B.A., Penrose D.L., Bernhardt E.S. 2011. Effects of urbanization and urban stream restoration on the physical and biological structure of stream ecosystems. Ecol. Appl. 21:1932–1949.

Walsh C.J., Imberger M., Burns M.J., Bos D.G., Fletcher T.D. 2022. Dispersed urban-stormwater control improved stream water quality in a catchment-scale experiment. Water Resour. Res. 58.

Walsh C.J., Roy A.H., Feminella J.W., Cottingham P.D., Groffman P.M., Morgan R.P. II. 2005. The urban stream syndrome: current knowledge and the search for a cure. J. North Am. Benthol. Soc. 24:706–723.

Walsh C.J., Webb J.A. 2016. Interactive effects of urban stormwater drainage, land clearance, and flow regime on stream macroinvertebrate assemblages across a large metropolitan region. Freshw. Sci. 35:324–339.

Wang S., Gao Y.-J., Wu D.-H., Xu D.-L., Wang T.-T., Fan S.-D., Wu E.-N., Song Y.-D., Zhang H.-J., Fu G.-P., Chen Z.-B., Mo L., Zhang Y., Ma Z.-L. 2024. Using a new fish indicator-based index with scoring and evaluation criteria to assess the ecological status in a disturbed subtropical river of China. Front. Ecol. Evol. 12.

Whitney J., Holloway J., Wright J., Boroughs K., Goodreau R., McManis A., Pistorius A., Puritty D., Ramirez M., Styers R. 2021. Assessing the invasion history and contemporary diet of nonnative redear sunfish (Lepomis microlophus Günther, 1859) in an ecotonal riverscape. Aquat. Invasions. 16:527–541.

Winemiller K.O., Rose K.A. 1992. Patterns of life-history diversification in North American fishes: Implications for population regulation. Can. J. Fish. Aquat. Sci. 49:2196–2218.

Yang B., Qu X., Liu H., Yang M., Xin W., Wang W., Chen Y. 2023. Urbanization reduces fish taxonomic and functional diversity while increases phylogenetic diversity in subtropical rivers. Sci Total Environ.:168178.

Zanaga D., Van De Kerchove R., Daems D., De Keersmaecker W., Brockmann C., Kirches G., Wevers J., Cartus O., Santoro M., Fritz S., Lesiv M., Herold M., Tsendbazar N.-E., Xu P., Ramoino F., Arino O. 2022. ESA WorldCover 10 m 2021 v200. .

Bourassa A.L., Fraser L., Beisner, B.E. 2017 Benthic macroinvertebrate and fish metacommunity structure in temperate urban streams. Journal of Urban Ecology.

