## Supplemental materials for "eDNA reveals urban habitat-specific sorting of a mixed regional fish fauna into distinct biodiversity and life-history assemblages"

**Supplemental Methods** … … … … … … … … … … … … … … … … … … .. 2

**Supplemental Figures** … … … … … … … … … … … … … … … … … … .. 6-17

**Supplemental Tables** … … … … … … … … … … … … … … … … … … .18-25

**References** … …. …. … … … … … … … … … … … … … … … … … … … … .26

### Supplemental Methods

#### *Taxonomic assignments*

Taxonomic identity decisions were informed by regional occurrence data for the Catawba and Yadkin–Pee Dee drainage basins, species accounts and trait information from FishBase, and regional fish distribution references for North Carolina. Decisions were intentionally conservative, prioritizing taxa already represented in the dataset to avoid artificially inflating species richness, thereby ensuring a coherent and defensible species-level dataset for downstream analyses. Specifically, ambiguous pairs were collapsed to a single species. Because some sequence reads could only be resolved to genus level, ambiguous detections were conservatively assigned to a single representative species for species-level analyses using a minimum diversity assignment rule to avoid inflating diversity estimates. For genera where two plausible candidate species occur in the region, assignments were made by directly comparing those species using known distributions, habitat associations, and sampling site characteristics. For example, *Nocomis leptocephalus* was selected instead of *N. micropogon* because all sampling sites were small Piedmont streams where *N. leptocephalus* commonly occurs, whereas *N. micropogon* is more typical of larger rivers. Similarly, *Erimyzon oblongus* was chosen over *E. sucetta* because the latter is more strongly associated with lentic wetlands, and *Ictalurus punctatus* was assigned instead of *I. furcatus* at sites located in stocked park ponds (e.g., Hornets Nest Park), where channel catfish are commonly stocked for recreational fishing. For *Micropterus* detections matching *M. salmoides/floridanus*, reads were assigned to the native lineage (*M. salmoides*). Additional pairwise comparisons were conducted for several taxa including *Ameiurus nebulosus* vs. *A. melas*, *Gambusia holbrooki* vs. *G. affinis*, and *Carpiodes cyprinus* vs. *C. carpio*, with assignments based on regional occurrence and compatibility with Piedmont stream habitats. Records assigned to *Moxostoma* were retained as *Moxostoma sp.* due to uncertainty consistent with an undescribed regional "Carolina Quillback" lineage that has known occurrences just outside of our sample area.

#### *Consistency of site-level eDNA detection signatures across field replicates and seasons*

To evaluate the consistency of site-level eDNA detection signatures, we analyzed fish detections at the level of individual field replicates. Analyses were conducted using presence/absence detections rather than read abundance to focus on reproducibility of taxon detection and to avoid treating metabarcoding read counts as direct estimates of organismal abundance, which can be affected by biological, sampling, amplification, and sequencing biases (Ficetola et al. 2015; Deagle et al. 2019). We first quantified replicate-level recurrence of taxon detections within each site-season combination. For each species detected at least once within a given site and season, we counted the number of field replicates in which that species was detected and classified detections as occurring in one, two, or three of the expected triplicate samples. These detection-frequency summaries were used to assess whether site-season detections were primarily represented by single-replicate events or by taxa recurring across multiple field replicates. This approach follows the broader use of biological and technical replication in eDNA metabarcoding to characterize detection reproducibility, improve taxon recovery, and reduce the influence of false-negative detections (Ficetola et al. 2015; Macher et al. 2021).

We next evaluated how observed site-level richness accumulated across replicate samples collected across both seasons for each site represented by two triplicate seasons (including zero-detection samples when no retained fish detections were present for a replicate). For each eligible site, replicate order was randomized 5000 times, and cumulative observed richness was calculated after each additional sample. The proportion of the full sample observed site fingerprint recovered after three samples was then summarized across randomizations. This analysis was used as a sample-based accumulation diagnostic to evaluate how sensitive observed site-level community fingerprints were to replicate sampling effort. This provided an expectation for the degree that additional field replication may increase taxon representation (Ficetola et al. 2015; Macher et al. 2021; Shirazi et al. 2021; Stauffer et al. 2021) and follows established guidance that accumulation metrics can be used to evaluate sampling dependence in observed richness, while avoiding overinterpretation of asymptotic richness when sample numbers are limited (Gotelli and Colwell 2001).

To test whether replicate eDNA samples retained site-level community structure, we calculated pairwise Jaccard dissimilarities among all replicate samples using the binary species matrix. Jaccard dissimilarity was used because it is an incidence-based measure appropriate for presence/absence community data and excludes shared absences from the similarity calculation. Pairwise comparisons were classified as within-site or between-site. The primary comparison was restricted to samples collected within the same season to control for seasonal differences in composition, and a second pooled comparison used all replicate samples across both seasons. For each comparison, we summarized the distribution of Jaccard dissimilarities and tested whether within-site dissimilarities were lower than between-site dissimilarities using permutation tests that randomized site labels while retaining the relevant season structure. This analysis directly evaluated whether replicate samples from the same site were more compositionally similar to one another than samples from different sites, consistent with eDNA studies that use incidence-based community dissimilarities to evaluate spatial structure and sampling reproducibility in detected assemblages (Macher et al. 2021; Courtaillac et al. 2024).

As a complementary multivariate test, we used permutational multivariate analysis of variance (PERMANOVA) to evaluate the relative contribution of site and season to variation in replicate-level community composition (Anderson 2001). PERMANOVA models were fit to the Jaccard dissimilarity matrix using site alone, season alone, and an additive season-plus-site model. These analyses were used as model-based support for the pairwise dissimilarity comparisons, with emphasis on variance explained by site identity rather than on PERMANOVA as the sole evidence for reproducibility. PERMANOVA was used because it provides a permutation-based framework for testing multivariate differences among groups from ecological distance matrices without requiring the distributional assumptions of classical multivariate analysis of variance (Anderson, 2001). All analyses were conducted in R using the vegan package for community dissimilarities and PERMANOVA (Oksanen et al. 2024).

*Biodiversity Responses to Urbanization-Associated Abiotic Variation*

To evaluate whether biodiversity indices tracked abiotic variables associated with urbanization, we compared site-level biodiversity metrics with measured and GIS-derived urbanization proxies. Raw biodiversity indices were merged with measured abiotic variables and GIS-derived landscape covariates by site. Biodiversity indices included species richness, Shannon diversity, Simpson diversity, mean Sørensen dissimilarity, Faith's phylogenetic diversity, mean pairwise phylogenetic distance, mean nearest taxon distance, functional richness, trait onion peeling, functional evenness, functional divergence, and Rao's quadratic entropy. Because several urbanization-associated variables were strongly redundant, we summarized the urbanization signal using a principal components analysis rather than fitting multivariable regressions with collinear predictors. The urbanization variable set included salinity, impervious cover within 100 m, impervious cover within 500 m, human-altered land cover within 500 m, developed land fraction within 1 km, corridor developed fraction, and the barrier-pressure index. These variables were z-scored prior to ordination, and the first principal component was retained as a shared site-level urbanization score. The sign of this axis was oriented so that larger values represented more urbanized conditions, and the resulting score was standardized before regression analysis. Pairwise Spearman correlations among the raw urbanization variables were calculated and visualized as a correlation heatmap to document predictor redundancy and to show the direction and magnitude of raw abiotic associations.

Each biodiversity index was then modeled separately as a function of the urbanization score using a one-predictor linear model of the form z(index) ~ z(urbanization score). Standardizing both predictor and response allowed slopes to be interpreted as standardized effect sizes and compared across indices measured on different scales. For each model, we extracted the standardized slope coefficient, 95% confidence interval, model p-value, and $R^2$. In parallel, Spearman rank correlations (Zar 2014) were calculated between each raw biodiversity index and the site-level urbanization score to evaluate whether monotonic relationships were consistent with the linear-model results. Benjamini–Hochberg correction (Benjamini and Hochberg 1995) was applied separately to the linear-model p-values and Spearman correlation p-values across biodiversity indices. Biodiversity indices were grouped for interpretation as retention-style metrics and labeled by biodiversity dimension as taxonomic, phylogenetic, or functional. Because Site 12 represented the only bounded-large site and occupied a distinct position in earlier environmental ordinations, the full analysis was repeated after excluding this site. All analyses were conducted in R using dplyr v.1.1.4 (Wickham 2023) for dataset integration and ggplot2 v3.5.1 for visualization (Wickham 2016).

### Supplemental Figures

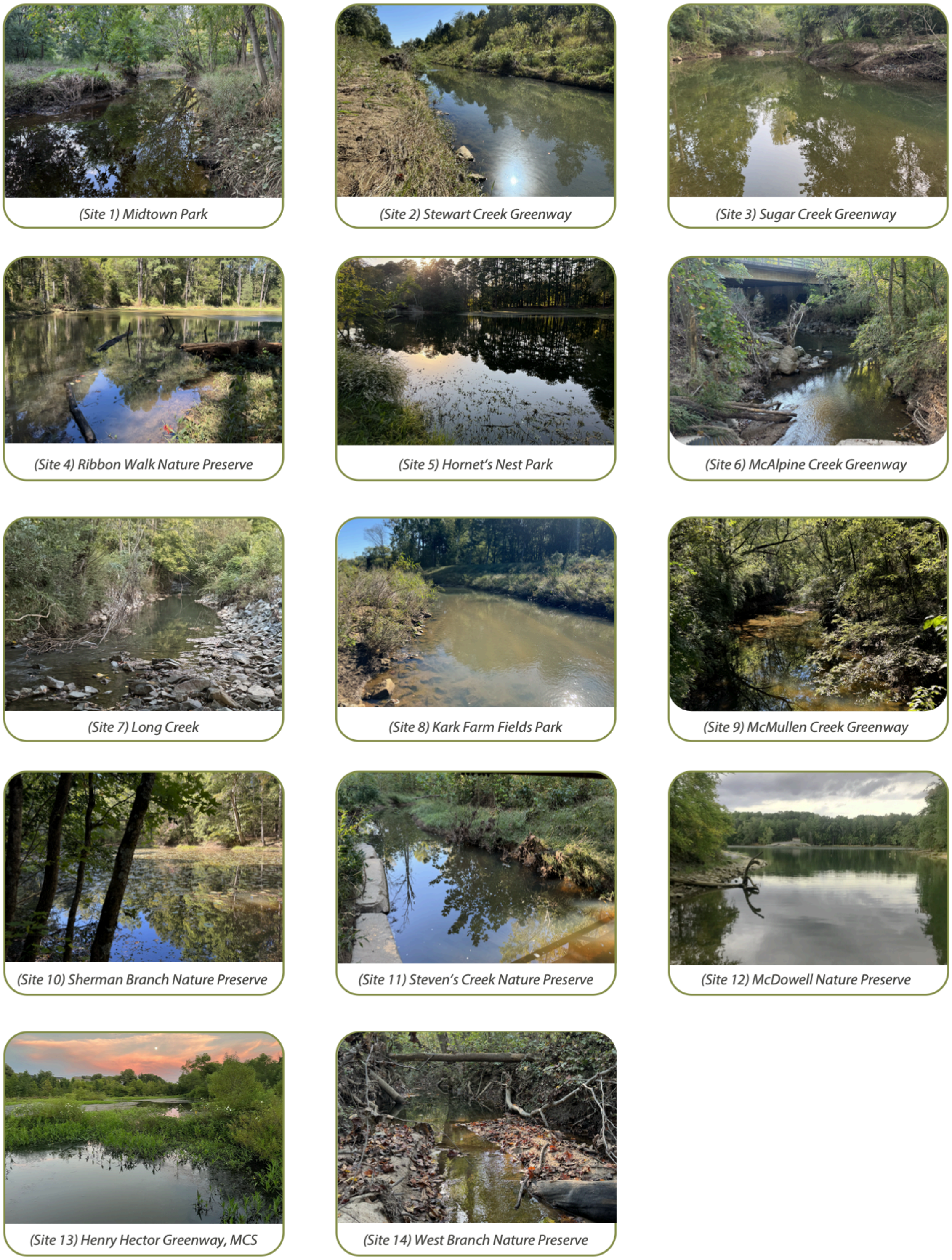


**Figure S1. Photos of sampling sites.** Site numbers correspond to site numbers used in the main text. All photos by KZ except Site 13 by AD.

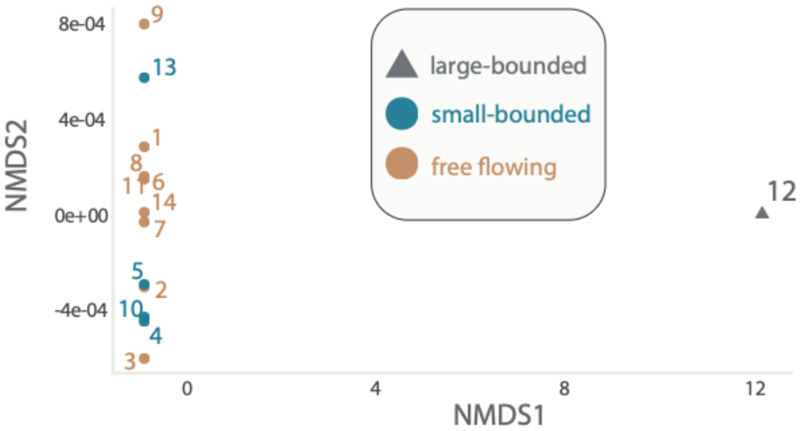


**Figure S2. Environmental NMDS based on z-scores from measured and GIS variables of all sites**. Points represent sites and shadings indicate habitat type with the triangle point indicating the large-bounded site.

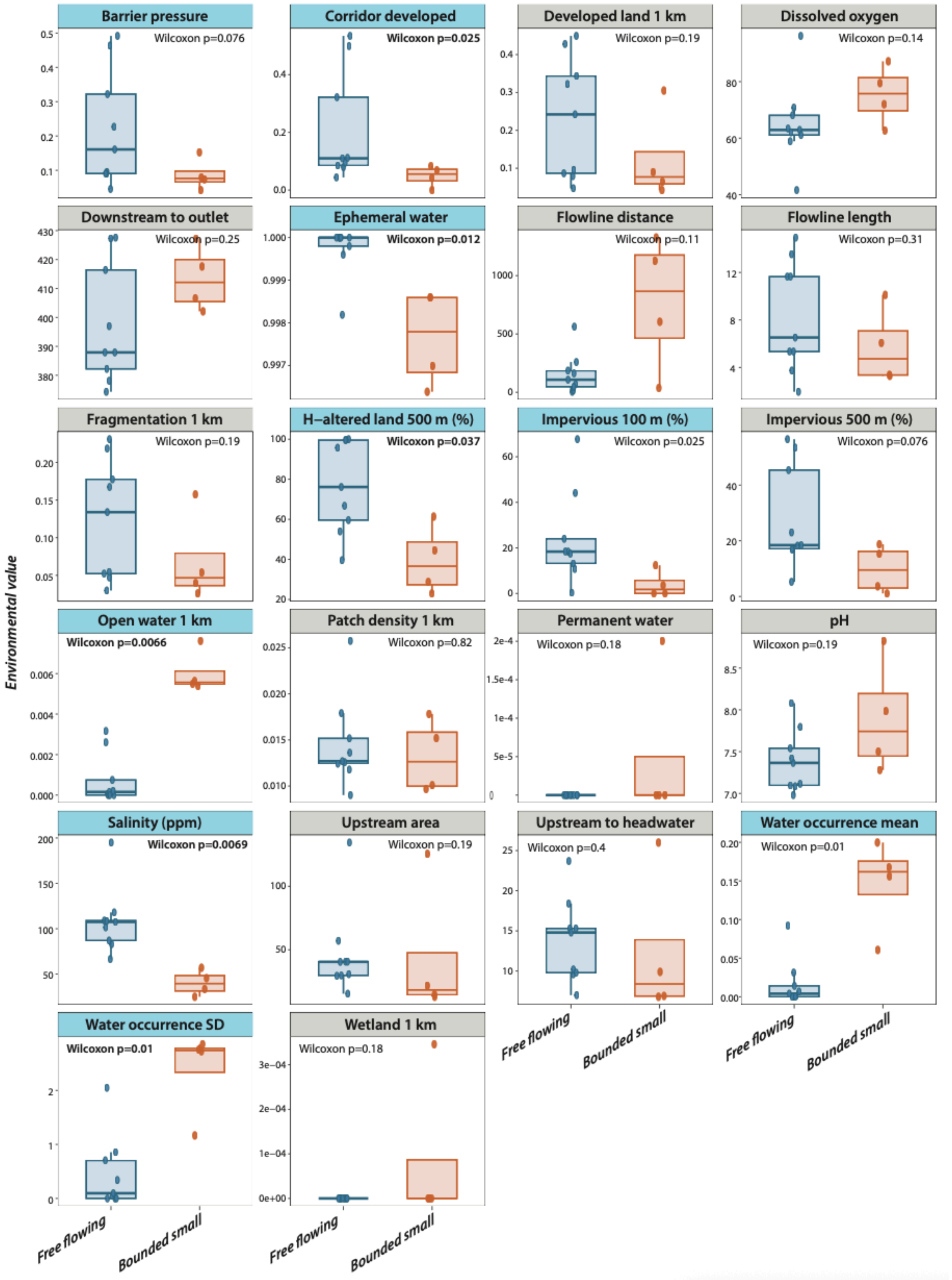


**Figure S3. Environmental Contrasts between bounded and free-flowing sites**. Boxplots depict the quantiles for the raw environmental values (Y axis) for each site type (x-axis) by variable (boxes). Points show individual values per site, shadings represent statistical significance based on the p-values corrected for multiple testing using the Benjamini-Hochberg false-discovery-rate procedure (Benjamini and Hochberg 1995) that are also indicated in each plot.

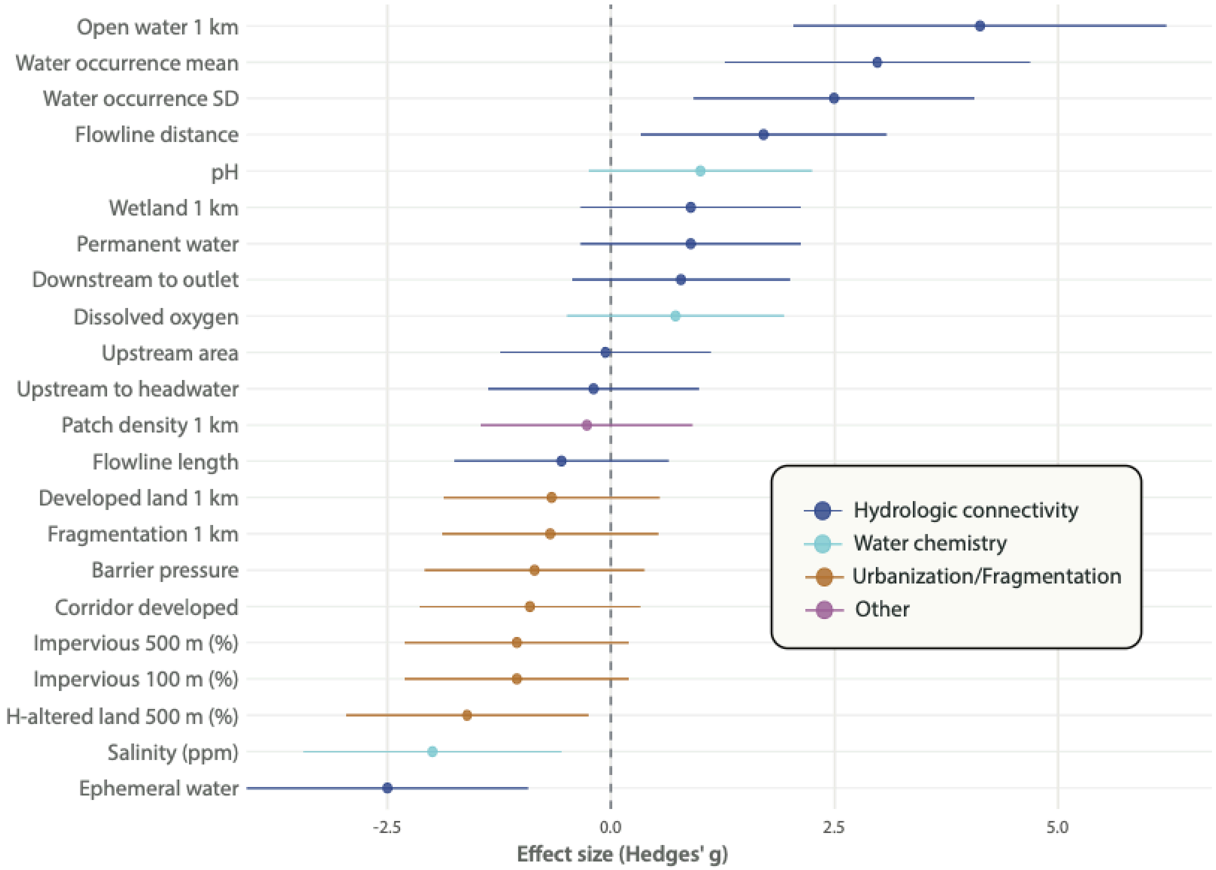


**Figure S4. Environmental contrasts between bounded small and free-flowing sites.** Hedges' g values depicting the standardized mean differences for environmental variables retained from the no-Site 12 environmental NMDS envfit analysis between small bounded and free flowing sites. Positive values indicate higher values in bounded small sites, and negative values indicate higher values in free-flowing sites. Error bars show approximate 95% confidence intervals. Effect sizes are intended as descriptive summaries of the environmental differences associated with habitat separation, not as independent tests of significance. H-altered = Human-altered.

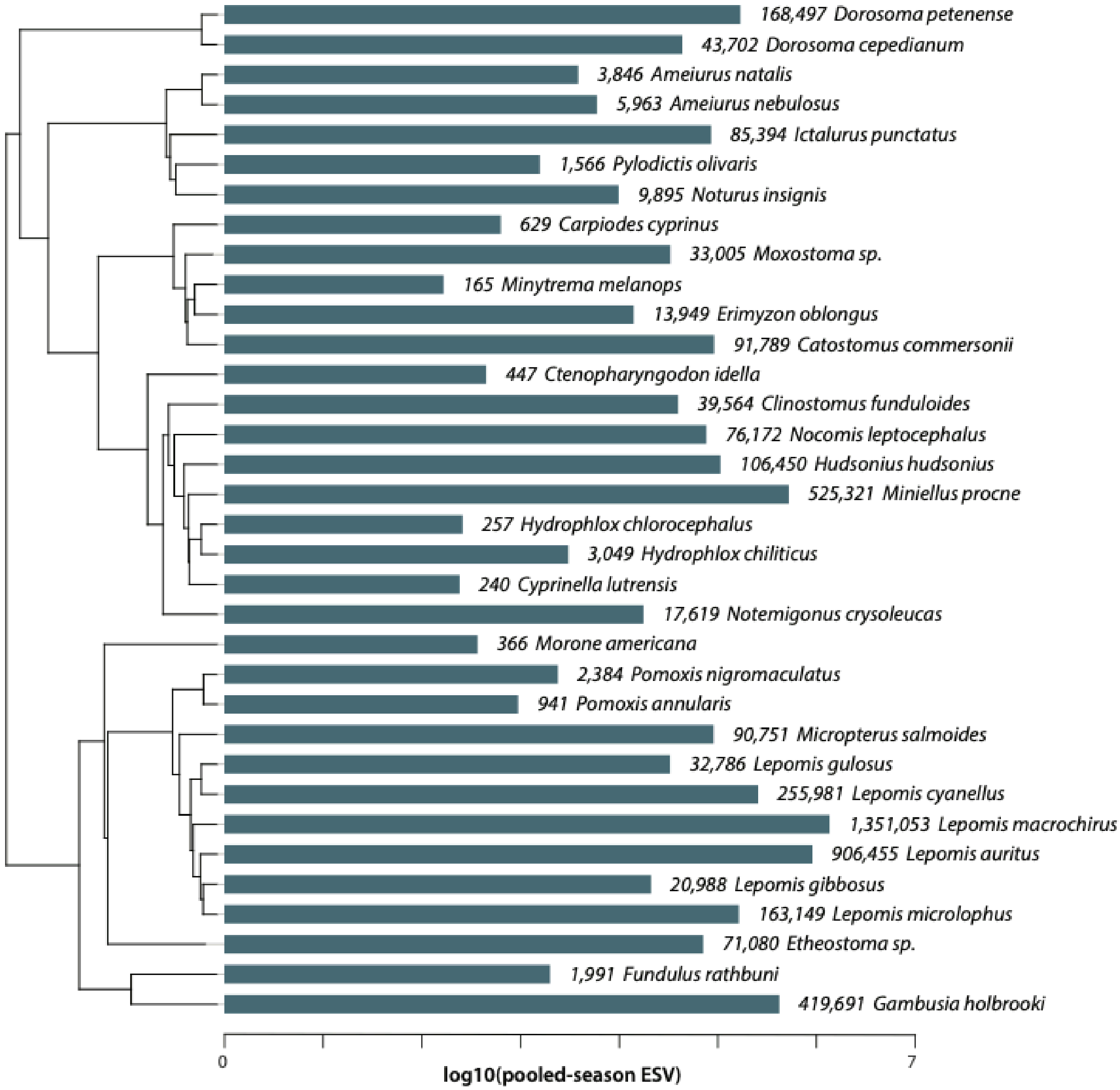


**Figure S5. ESV abundance across detected fish taxa.** Horizontal bars show pooled-season environmental sequence variant (ESV) abundance for each detected fish taxon, plotted on a log10 scale. Taxa are ordered according to their evolutionary relationships depicted by the phylogenetic tree (left). Numeric labels beside each bar give the raw pooled-season ESV total for that taxon, followed by the taxon name.

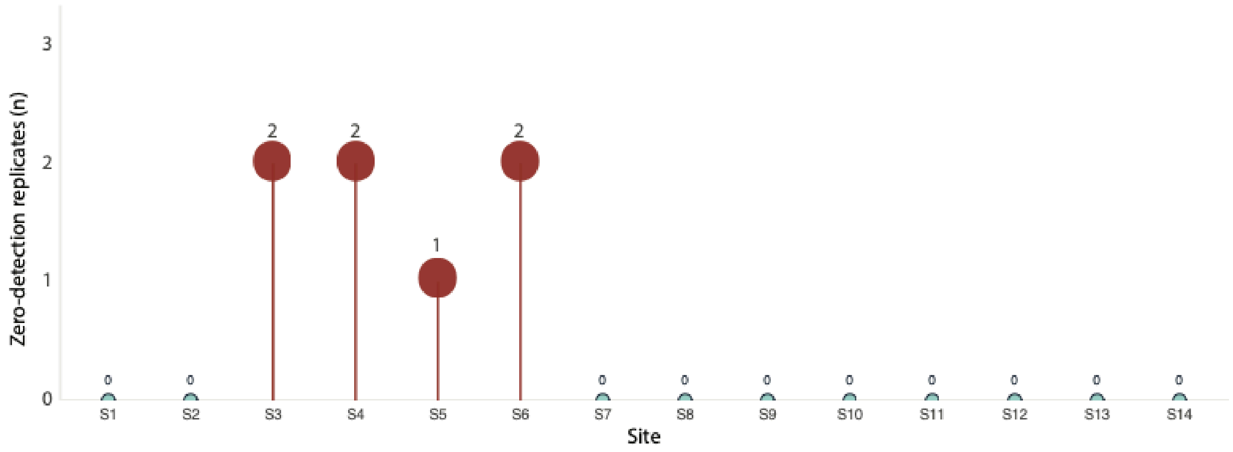


**Figure S6. Zero-detection biological replicates by site.** Points show the number of replicate samples that contained zero retained fish detections. Vertical lollipop stems connect each site to zero to emphasize the count scale; numeric labels give the exact number of replicates with zero-detections per site.

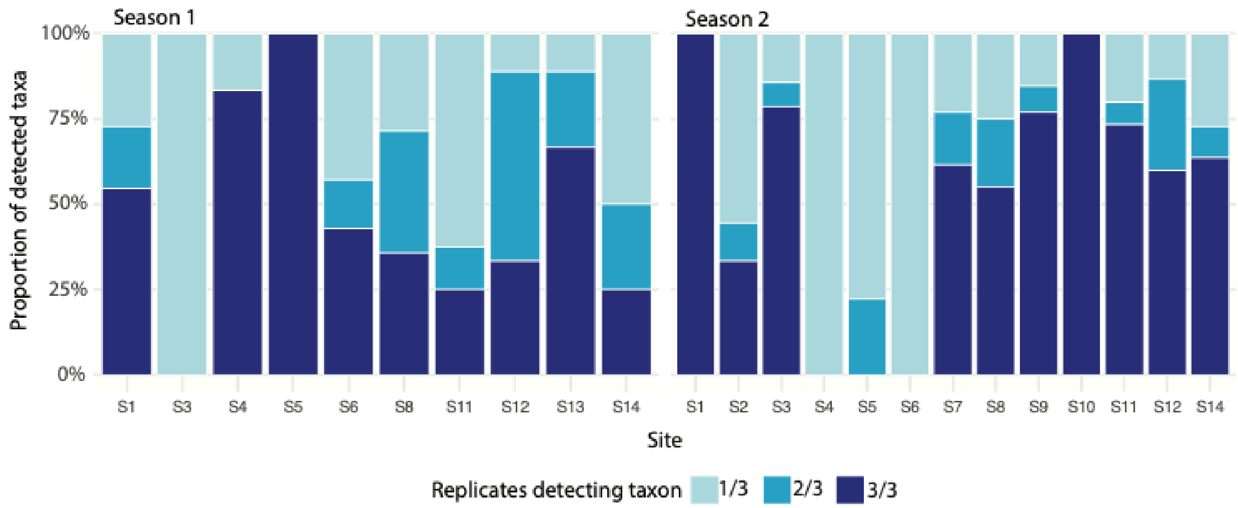


**Figure S7. Replicate detection frequency across site-season combinations**. Each bar represents one sampled site within a season, and stacked colors show the proportion of detected taxa recovered in one, two, or all three expected field replicates. Facets separate sampling seasons; only site-season combinations with at least one retained fish detection are shown.

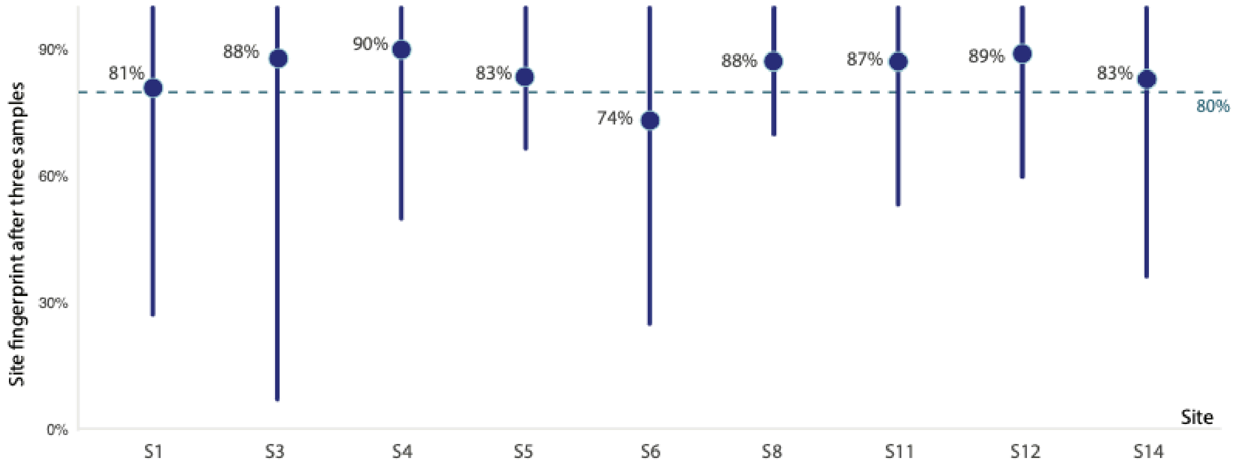


**Figure S8. Randomized site-level accumulation of observed fish community fingerprints.** Points show the mean percentage of each site's full observed six-sample richness recovered after three randomly ordered replicate samples across 5000 randomizations. Vertical intervals show the 95% random-order interval, and the dashed horizontal line marks 80% recovery. Only sites with two reconstructed triplicate seasons are included.

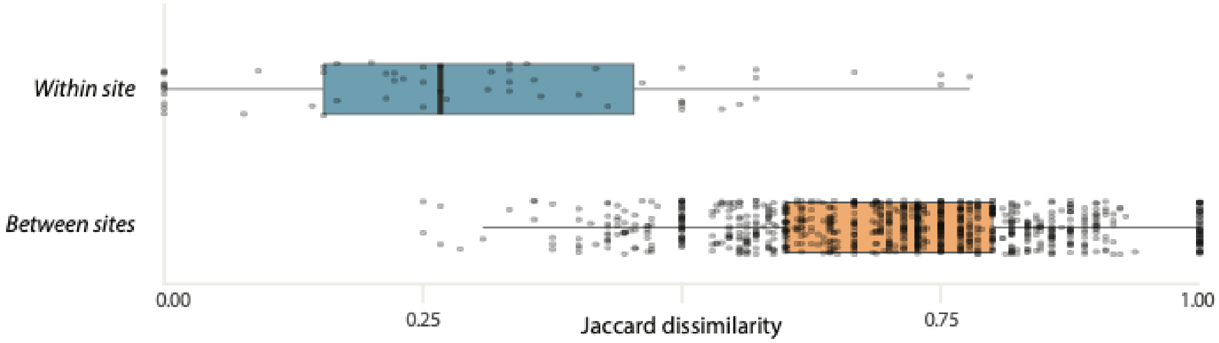


**Figure S9. Season-controlled pairwise Jaccard dissimilarity among replicate fish eDNA samples**. Points represent pairwise comparisons among replicate samples collected in the same season. Horizontal boxplots summarize the distribution of Jaccard dissimilarity for within-site same-season pairs and between-site same-season pairs; lower values indicate more similar presence/absence composition.

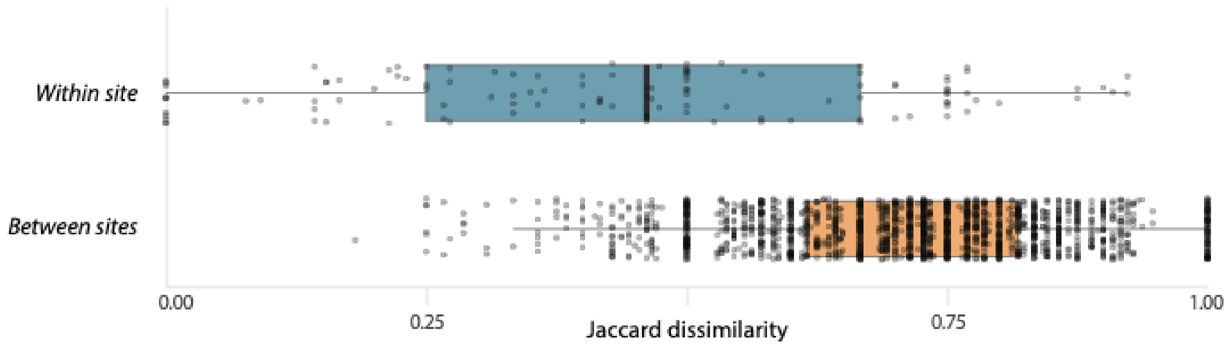


**Figure S10. Pooled pairwise Jaccard dissimilarity among replicate fish eDNA samples.** Points represent all pairwise comparisons among replicate samples across the full data set. Horizontal boxplots summarize the distribution of Jaccard dissimilarity for pairs from the same site and pairs from different sites; lower values indicate more similar presence/absence composition.

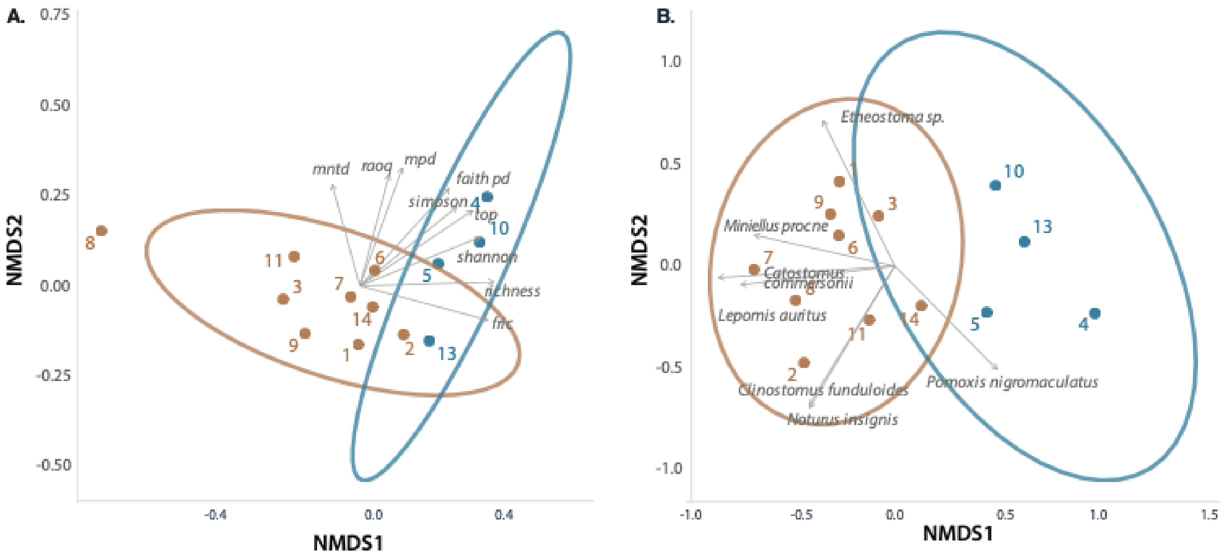


**Figure S11. NMDS ordinations recalculated after excluding Site 12, the single bounded-large site.** **(A)** Diversity-index NMDS for the remaining 13 sites (Stress = 0.030 | PERMANOVA p = 0.005). **(B)** Species presence-absence NMDS for the same reduced site set (Stress = 0.117 | PERMANOVA p = 0.003).

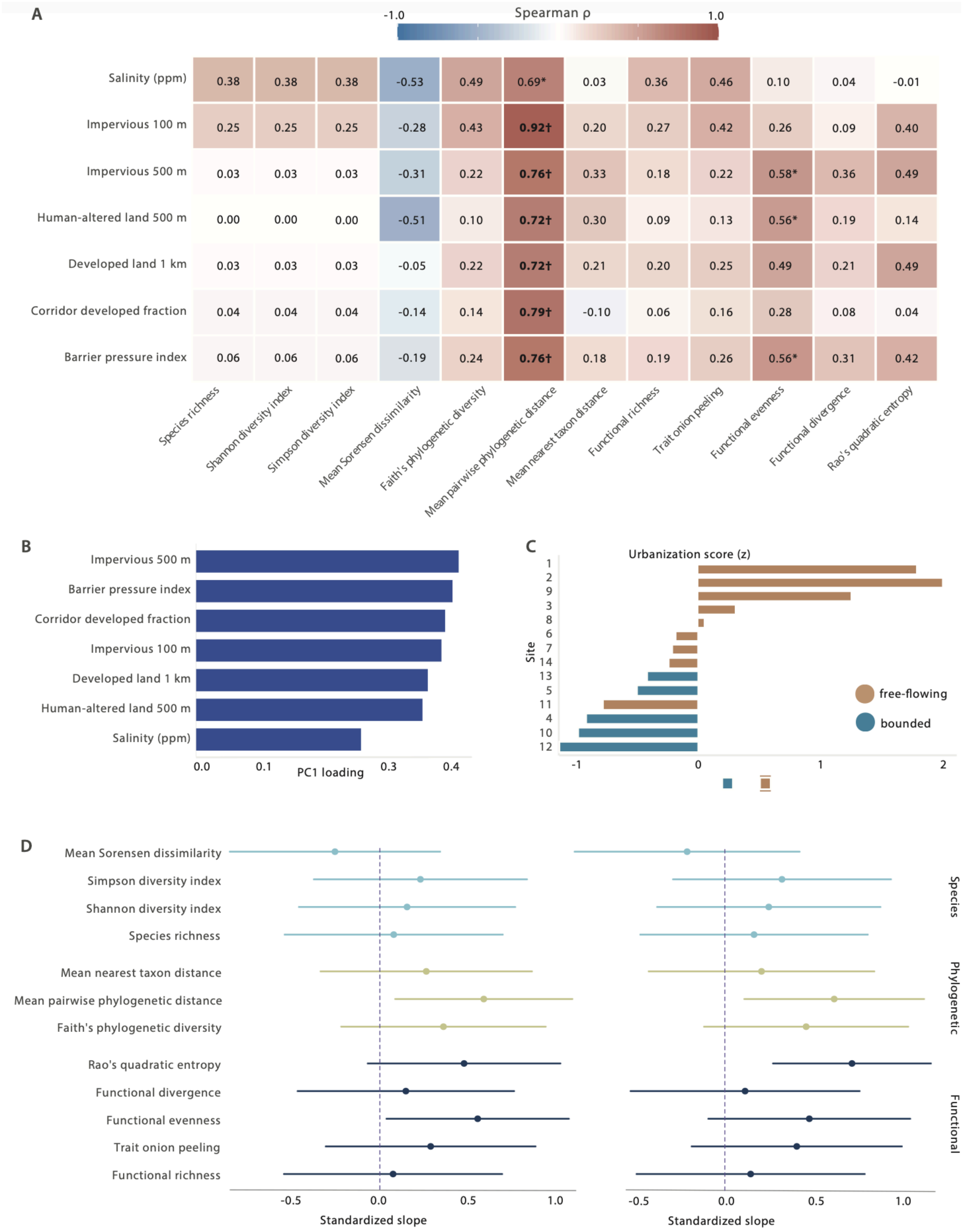


**Figure S12. Associations between urbanization-associated abiotic variables and multidimensional biodiversity indices across sampled sites. (A)** Heat map of raw pairwise Spearman correlations between abiotic variables and biodiversity indices. Cells show Spearman correlations with cell shading corresponding to direction (warm:positive & cool:negative). * indicates $p < 0.05$ and † indicates significance after Benjamini–Hochberg correction ($p < 0.05$). **(B)** Loadings of the seven focal abiotic variables on the first principal component (PC1) that captured 78% of the total variation and used to summarize the shared urbanization gradient. Higher loadings indicate stronger contribution to the composite urbanization score. **(C)** Site-level values of the urbanization score (z-standardized PC1), ordered from negative to positive. Shadings correspond to habitat category shadings in the main text, with warmer shading for free-flowing sites and cooler shading for bounded sites. **(D)** Standardized slopes and 95% confidence intervals from linear models relating each biodiversity index to the urbanization score for **(left)** all 14 sites, and **(right)** the sensitivity analysis excluding Site 12. The vertical dashed line marks zero effect. Indices are grouped by biodiversity dimension (species, phylogenetic, and functional).

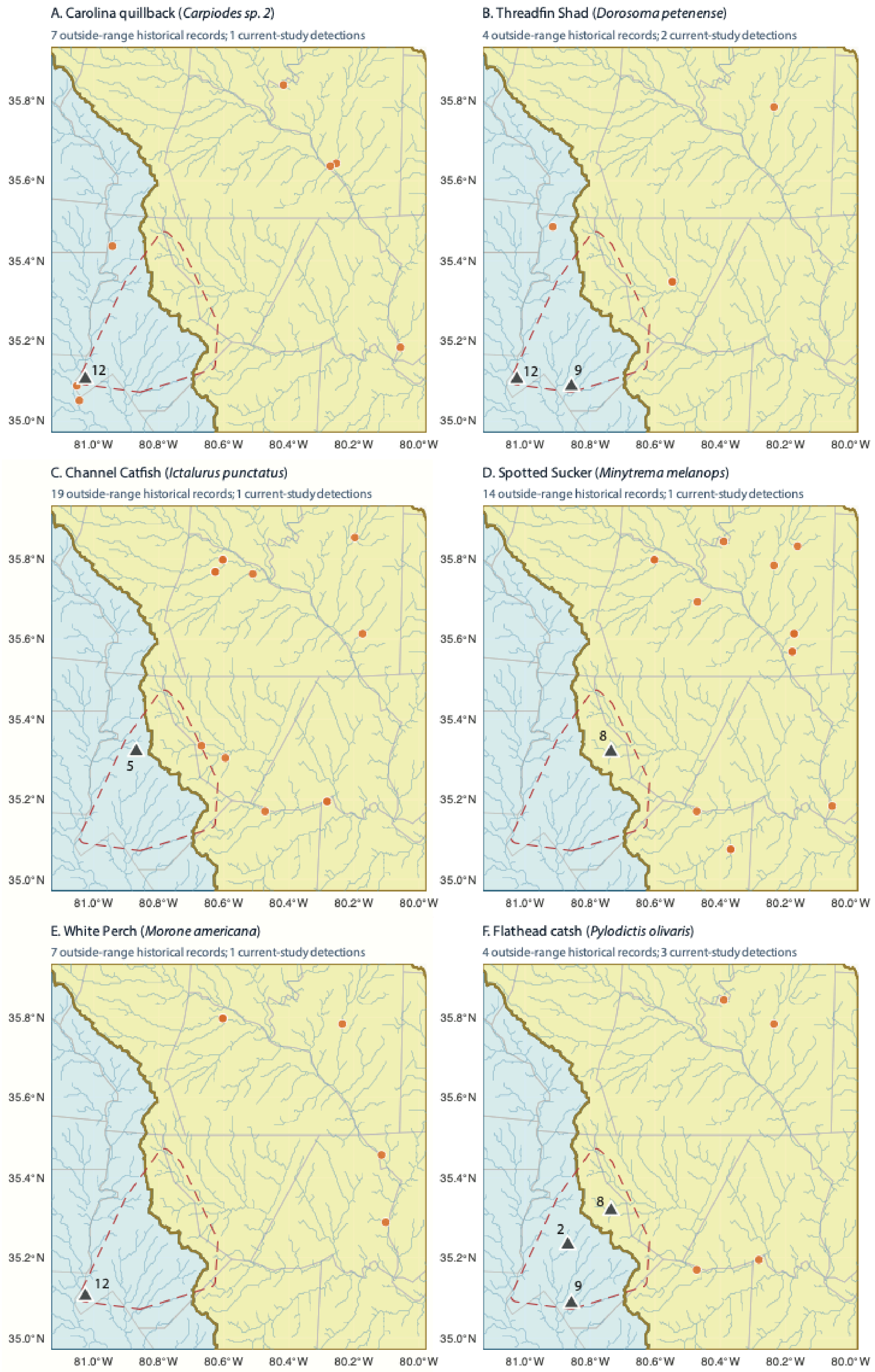
A. Carolina quillback (*Carpiodes sp. 2*)
7 outside-range historical records; 1 current-study detections
12
B. Threadfin Shad (*Dorosoma petenense*)
4 outside-range historical records; 2 current-study detections
12
9
C. Channel Catfish (*Ictalurus punctatus*)
19 outside-range historical records; 1 current-study detections
5
D. Spotted Sucker (*Minytrema melanops*)
14 outside-range historical records; 1 current-study detections
8
E. White Perch (*Morone americana*)
7 outside-range historical records; 1 current-study detections
12
F. Flathead catsh (*Pylodictis olivaris*)
4 outside-range historical records; 3 current-study detections
8
2
9
35.8°N
35.6°N
35.4°N
35.2°N
35.0°N
81.0°W
80.8°W
80.6°W
80.4°W
80.2°W
80.0°W

**Figure S13. Nearest historic records of taxa without prior public records in the Charlotte metropolitan area of focus in this study.** (A) Carolina Quillback; (B) Threadfin Shad; (C) Channel Catfish; (D) Spotted Sucker; (E) White Perch; (F) Flathead Catfish. Records are taken from the NC Freshwater fish archive (LeGrand, H., J. Amoroso, and T. Howard. 2026) which draws from museum and agency records and is consistent with the records utilized in the NC freshwater fish atlas (Tracy et al. 2020)**.** Each panel shows curated outside-range historical records as orange circles, current-study detections as gray triangles labeled by site number, the buffered study footprint as a dashed red polygon, HydroRIVERS flowlines in blue, and the visible portions of the Catawba and Yadkin-Pee Dee drainage basins as pale background polygons.

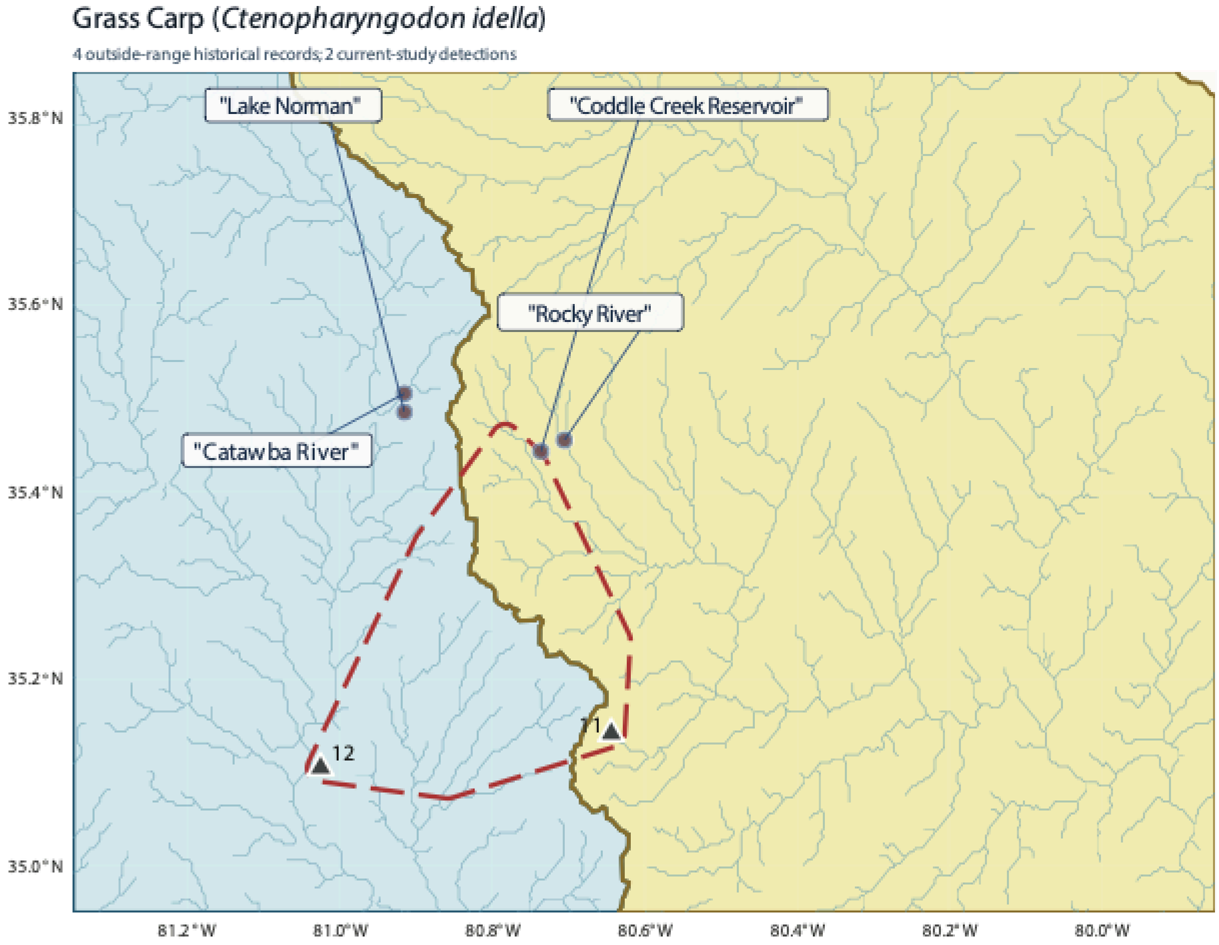


**Figure S14. Grass Carp regional context map.** Gray triangles indicate study detections and labeled points indicate named rivers and reservoirs supplied from the referenced USGS NAS locality notes. Named localities are in quotes to represent uncertainty as the available USGS NAS records contained no coordinates or other metadata besides the county and name of the water body. As such, the plotted Grass Carp anchors are approximate map positions intended to indicate the named water bodies rather than exact collection coordinates. North Carolina also permits stocking of Grass Carp in private ponds, so the mapped localities should be interpreted as documented examples rather than the full extent of occurrence.

### Supplemental Tables

**Table S1. PERMANOVA summary for the impact of site identity and season on eDNA detection replicates**

| model | term | Df | SS | $R^2$ | F | Pr(>F) |
|---|---|---|---|---|---|---|
| jaccard_dist ~ site | Model | 13 | 9.85 | **0.62** | 6.25 | **0.001** |
| jaccard_dist ~ site | Residual | 48 | 5.82 | 0.37 | — | — |
| jaccard_dist ~ site | Total | 61 | 15.67 | 1 | — | — |
| jaccard_dist ~ season | Model | 1 | 0.50 | **0.03** | 1.97 | **0.042** |
| jaccard_dist ~ season | Residual | 60 | 15.17 | 0.97 | — | — |
| jaccard_dist ~ season | Total | 61 | 15.67 | 1 | — | — |
| jaccard_dist ~ season + site | Model | 14 | 10.19 | **0.65** | 6.24 | **0.001** |
| jaccard_dist ~ season + site | Residual | 47 | 5.48 | 0.35 | — | — |
| jaccard_dist ~ season + site | Total | 61 | 15.67 | 1 | — | — |

**Table S2. Ecological and Life History Data**

| | Feeding Ecology | | | | | Water Column Usage | | | | | Reproduction | | |
|---|---|---|---|---|---|---|---|---|---|---|---|---|---|
| Taxa | Pisc. | Inv. | Pla. | Her. | Detr. | Ben. | Dem. | Pel. | Surf. | TL | Eggs | R.A. | Ref. |
| *Ameiurus natalis* | 2 | 4 | 0 | 4 | 3 | 5 | 4 | 0 | 0 | 47 | 5000 | 3 | (Rohde et al. 1995) |
| *Ameiurus nebulosus* | 4 | 4 | 0 | 3 | 2 | 5 | 4 | 1 | 0 | 53.2 | 13000 | 2 | (Rohde et al. 1995) |
| *Carpiodes sp2.* | 0 | 3 | 0 | 1 | 5 | 5 | 4 | 1 | 0 | 66 | 360000 | 6 | (Parker and Franzin 1991; Lamprecht et al. 2015; IDNR 2024) |
| *Catostomus commersonii* | 0 | 4 | 1 | 4 | 3 | 4 | 5 | 0 | 0 | 63.5 | 50000 | 4 | (Rohde et al. 1995) |
| *Clinostomus funduloides* | 0 | 5 | 0 | 0 | 0 | 1 | 2 | 4 | 2 | 10.9 | 800 | 1 | (Rohde et al. 1995) |
| *Ctenopharyngodon idella* | 0 | 1 | 0 | 5 | 3 | 3 | 3 | 3 | 3 | 125 | 500000 | 3 | (Rohde et al. 1995; Wilson et al. 2025) |
| *Cyprinella lutrensis* | 1 | 4 | 0 | 0 | 0 | 3 | 3 | 3 | 3 | 9 | 212800 | 2 | (Rohde et al. 1995; Nico et al. 2025) |
| *Dorosoma cepedianum* | 0 | 1 | 5 | 2 | 0 | 0 | 0 | 3 | 5 | 52 | 378,900 | 2 | (Bodola 1966; Shaw 2023) |

| | | | | | | | | | | | | | |
|---|---|---|---|---|---|---|---|---|---|---|---|---|---|
| *Dorosoma petenense* | 0 | 1 | 5 | 2 | 0 | 0 | 0 | 3 | 5 | 52 | 20000 | 1 | (Pauly 2010; Hammerson 2011) |
| *Erimyzon oblongus* | 0 | 4 | 3 | 4 | 1 | 5 | 4 | 0 | 0 | 36 | 72000 | 2 | (Rohde et al. 1995) |
| *Etheostoma sp.** | 0 | 5 | 0 | 0 | 0 | 3 | 3 | 3 | 2 | 7.2 | 500 | 1 | (Rohde et al. 1995) |
| *Fundulus diaphanus* | 0 | 5 | 1 | 3 | 2 | 0 | 0 | 2 | 3 | 13 | 250 | 2 | (Rohde et al. 1995) |
| *Gambusia holbrooki* | 0 | 5 | 1 | 0 | 0 | 0 | 0 | 2 | 5 | 6.5 | 315 | 1 | (Rohde et al. 1995) |
| *Ictalurus punctatus* | 4 | 4 | 0 | 3 | 1 | 5 | 4 | 0 | 0 | 127 | 21000 | 4 | (Rohde et al. 1995) |
| *Lepomis auritus* | 3 | 4 | 0 | 0 | 0 | 2 | 4 | 2 | 1 | 24 | 14000 | 3 | (Rohde et al. 1995) |
| *Lepomis cyanellus* | 4 | 4 | 0 | 0 | 0 | 1 | 4 | 3 | 1 | 31 | 10000 | 1 | (Rohde et al. 1995; Fofonoff et al. 2018) |
| *Lepomis gibbosus* | 4 | 5 | 0 | 0 | 0 | 1 | 4 | 4 | 1 | 38.1 | 3000 | 2 | (Rohde et al. 1995) |
| *Lepomis gulosus* | 4 | 4 | 0 | 0 | 0 | 2 | 5 | 1 | 0 | 31 | 63000 | 1 | (Rohde et al. 1995) |
| *Lepomis macrochirus* | 4 | 4 | 0 | 0 | 0 | 4 | 4 | 4 | 1 | 41 | 180000 | 1 | (Rohde et al. 1995) |
| *Lepomis microlophus* | 0 | 5 | 0 | 0 | 0 | 4 | 3 | 0 | 0 | 38.1 | 45000 | 2 | (Rohde et al. 1995) |
| *Micropterus salmoides* | 5 | 2 | 0 | 0 | 0 | 1 | 4 | 3 | 1 | 97 | 100000 | 2 | (Rohde et al. |

| | | | | | | | | | | | | | |
|---|---|---|---|---|---|---|---|---|---|---|---|---|---|
| | | | | | | | | | | | 0 | | 1995) |
| *Minytrema melanops* | 0 | 5 | 0 | 0 | 0 | 4 | 4 | 0 | 0 | 49.5 | 129732 | 3 | (Rohde et al. 1995; Agbugui, M. O., & Adeniyi, A. O. 2021) |
| *Morone americana* | 3 | 3 | 0 | 0 | 0 | 0 | 2 | 4 | 1 | 48.3 | 457000 | 4 | (Rohde et al. 1995; U.S. Geological Survey 2026) |
| *Moxostoma erythrurum* | 0 | 4 | 0 | 3 | 4 | 4 | 4 | 0 | 0 | 51 | 5000 | 4 | (Rohde et al. 1995) |
| *Nocomis leptocephalus* | 1 | 4 | 0 | 4 | 1 | 4 | 4 | 1 | 0 | 26 | 800 | 2 | (Rohde et al. 1995) |
| *Notemigonus crysoleucas* | 0 | 4 | 2 | 4 | 0 | 0 | 0 | 4 | 4 | 30 | 4000 | 1 | (Rohde et al. 1995) |
| *Hydrophlox chiliticus* | 0 | 3 | 2 | 3 | 0 | 1 | 3 | 3 | 1 | 6.5 | 5000 | 1 | (iNaturalist 2026a) |
| *Hydrophlox chlorocephalus* | 0 | 3 | 2 | 3 | 0 | 1 | 3 | 3 | 1 | 7.6 | 5000 | 1 | (Pauly 2010; iNaturalist 2026b) |
| *Hudsonius hudsonius* | 0 | 4 | 2 | 3 | 0 | 0 | 0 | 4 | 1 | 8.2 | 5000 | 1 | (Rohde et al. 1995) |
| *Miniellus procne* | 0 | 4 | 2 | 3 | 0 | 1 | 4 | 2 | 0 | 7.2 | 2500 | 2 | (Frimpong and Angermeier 2009; Lagacy 2024) |

| | | | | | | | | | | | | | |
|---|---|---|---|---|---|---|---|---|---|---|---|---|---|
| *Noturus insignis* | 2 | 4 | 0 | 0 | 3 | 4 | 4 | 0 | 0 | 15 | 107 | 1 | (Rohde et al. 1995) |
| *Pomoxis annularis* | 3 | 3 | 2 | 0 | 0 | 2 | 4 | 3 | 0 | 53 | 213000 | 2 | (Rohde et al. 1995) |
| *Pomoxis nigromaculatus* | 5 | 2 | 2 | 0 | 0 | 1 | 4 | 4 | 1 | 49 | 188000 | 2 | (Rohde et al. 1995; NJFW 2026) |
| *Pylodictis olivaris* | 5 | 2 | 0 | 0 | 0 | 5 | 3 | 0 | 0 | 155 | 100000 | 4 | (Rohde et al. 1995) |

*Pisc = Piscivory; Inv = Invertivory; Pla = Planktivory; Her = Herbivory; Det = Detritivory; Ben = Benthic; Dem = Demersal; Pel = Pelagic; Sur = Surface; TL = Maximum Total Length (cm); Eggs = eggs per year (upper bound); R.A. = Reproductive age (lower bound, years)* .
*Etheostoma sp. *based on* E. olmstedi *which previously included the local species* E. maculaticeps.

**Table S3. Diversity Index Values by Site**

| Site | Species richness | Shannon | Simpson | Sorensen (mean) | Faith's PD | Phylo MPD | Phylo MNTD | Fric | Top | Feve | Fdiv | Raoq | Habitat type |
|---|---|---|---|---|---|---|---|---|---|---|---|---|---|
| 1 | 11 | 2.4 | 0.91 | 0.45 | 1077.53 | 290.93 | 126.78 | 8.61 | 18.39 | 0.88 | 0.79 | 9.78 | free flowing |
| 2 | 9 | 2.2 | 0.89 | 0.53 | 854.55 | 280.59 | 100.74 | 6.19 | 17.12 | 0.92 | 0.77 | 9.62 | free flowing |
| 3 | 14 | 2.64 | 0.93 | 0.41 | 1124.13 | 265.56 | 104.84 | 17.65 | 26.1 | 0.86 | 0.78 | 8.79 | free flowing |
| 6 | 12 | 2.48 | 0.92 | 0.44 | 965.88 | 266.48 | 108.35 | 2.09 | 16.12 | 0.86 | 0.8 | 6.36 | free flowing |
| 7 | 13 | 2.56 | 0.92 | 0.49 | 1118.46 | 280.22 | 111.34 | 6.54 | 18.89 | 0.76 | 0.76 | 7.99 | free flowing |
| 8 | 20 | 3 | 0.95 | 0.48 | 1297.4 | 268.97 | 74.78 | 42.93 | 33.45 | 0.86 | 0.79 | 9.11 | free flowing |
| 9 | 13 | 2.56 | 0.92 | 0.46 | 1123.37 | 273.25 | 124.69 | 15.05 | 24.61 | 0.84 | 0.75 | 9.55 | free flowing |
| 11 | 15 | 2.71 | 0.93 | 0.43 | 999.53 | 256.55 | 79.84 | 19.74 | 21.41 | 0.86 | 0.79 | 7.78 | free flowing |
| 14 | 11 | 2.4 | 0.91 | 0.45 | 914.02 | 258.89 | 103.18 | 7.03 | 16.91 | 0.89 | 0.81 | 7.85 | free flowing |
| 4 | 6 | 1.79 | 0.83 | 0.61 | 394.42 | 120.02 | 70.39 | 0.02 | 4.15 | 0.78 | 0.76 | 6.77 | bounded small |
| 5 | 9 | 2.2 | 0.89 | 0.47 | 738.95 | 208.82 | 120.06 | 2.5 | 8.33 | 0.84 | 0.75 | 7.11 | bounded small |
| 10 | 6 | 1.79 | 0.83 | 0.53 | 545.28 | 201.57 | 129.33 | 0.12 | 5.29 | 0.85 | 0.73 | 6.39 | bounded small |
| 12 | 15 | 2.71 | 0.93 | 0.52 | 1134.49 | 250.92 | 92.07 | 19.57 | 26.14 | 0.79 | 0.77 | 10.55 | bounded large |
| 13 | 9 | 2.2 | 0.89 | 0.5 | 866.56 | 249.92 | 147.77 | 5.23 | 15.69 | 0.87 | 0.81 | 9.86 | bounded small |

Sites correspond to site numbers in the main text. PD=Phylogenetic diversity; MPD = mean pairwise phylogenetic distance; MNTD = mean nearest taxon distance; Fric = functional richness; Top = trait onion peeling area; Feve = functional evenness; Fdiv = functional divergence; Raoq = Rao's quadratic entropy.

**Table S4. Associations between the urbanization score and biodiversity indices across all 14 sites**

| Index | Dimension | Spearman rho | BH-adjusted Spearman p | Standardized slope | 95% CI |
|---|---|---|---|---|---|
| **Mean pairwise phylogenetic distance** | **Phylogenetic** | **0.873** | **0.00056** | **0.586** | **[0.076, 1.096]** |
| Functional evenness | Functional | 0.486 | 0.46966 | 0.551 | [0.026, 1.076] |
| Mean Sorensen dissimilarity | Species | -0.345 | 0.51975 | -0.271 | [-0.876, 0.335] |
| Trait onion peeling | Functional | 0.341 | 0.51975 | 0.280 | [-0.324, 0.884] |
| Faith’s phylogenetic diversity | Phylogenetic | 0.336 | 0.51975 | 0.354 | [-0.235, 0.942] |
| Rao’s quadratic entropy | Functional | 0.323 | 0.51975 | 0.472 | [-0.082, 1.027] |
| Functional divergence | Functional | 0.257 | 0.56468 | 0.137 | [-0.486, 0.760] |
| Functional richness | Functional | 0.253 | 0.56468 | 0.064 | [-0.564, 0.691] |
| Mean nearest taxon distance | Phylogenetic | 0.174 | 0.56468 | 0.255 | [-0.353, 0.863] |
| Species richness | Species | 0.169 | 0.56468 | 0.068 | [-0.560, 0.695] |
| Shannon diversity index | Species | 0.169 | 0.56468 | 0.144 | [-0.478, 0.767] |
| Simpson diversity index | Species | 0.169 | 0.56468 | 0.221 | [-0.392, 0.835] |

**Table S5. Associations between the urbanization score and biodiversity indices after excluding Site 12**

| Index | Dimension | Spearman rho | BH-adjusted Spearman p | Standardized slope | 95% CI |
|---|---|---|---|---|---|
| **Mean pairwise phylogenetic distance** | **Phylogenetic** | **0.907** | **0.00023** | **0.626** | **[0.109, 1.144]** |
| Rao's quadratic entropy | Functional | 0.654 | 0.07061 | 0.729 | [0.274, 1.183] |
| Trait onion peeling | Functional | 0.626 | 0.07061 | 0.411 | [-0.194, 1.016] |
| Faith's phylogenetic diversity | Phylogenetic | 0.621 | 0.07061 | 0.465 | [-0.123, 1.052] |
| Functional richness | Functional | 0.500 | 0.19647 | 0.145 | [-0.511, 0.802] |
| Functional evenness | Functional | 0.407 | 0.22640 | 0.483 | [-0.098, 1.064] |
| Species richness | Species | 0.405 | 0.22640 | 0.165 | [-0.490, 0.819] |
| Shannon diversity index | Species | 0.405 | 0.22640 | 0.250 | [-0.393, 0.892] |
| Simpson diversity index | Species | 0.405 | 0.22640 | 0.326 | [-0.302, 0.953] |
| Functional divergence | Functional | 0.275 | 0.41884 | 0.113 | [-0.546, 0.773] |
| Mean Sorensen dissimilarity | Species | -0.264 | 0.41884 | -0.220 | [-0.868, 0.427] |
| Mean nearest taxon distance | Phylogenetic | 0.044 | 0.88662 | 0.208 | [-0.441, 0.857] |

### References

Agbugui, M. O., & Adeniyi, A. O. 2021. biology spotted sucker Minytrema melanops Rafinesque, 1820 from River Niger Agenebode. Int. J. Fish. Aquat. Res. 6:82–90.

Anderson M.J. 2001. A new method for non‐parametric multivariate analysis of variance. Austral Ecol. 26:32–46.

Benjamini Y., Hochberg Y. 1995. Controlling the false discovery rate: A practical and powerful approach to multiple testing. J. R. Stat. Soc. Series B Stat. Methodol. 57:289–300.

Bodola A. 1966. Life history of the gizzard shad, Dorosoma cepedianum (Le Sueur), in western Lake Erie. Available from https://www.usgs.gov/publications/life-history-gizzard-shad-dorosoma-cepedianum-le-sueur-western-lake-erie.

Courtaillac K.-L., Landschoff J., Hull K., von der Heyden S. 2024. The effect of spatio‐temporal sampling and biological replication on the detection of kelp forest fish communities using eDNA metabarcoding. Environ. DNA. 6.

Deagle B.E., Thomas A.C., McInnes J.C., Clarke L.J., Vesterinen E.J., Clare E.L., Kartzinel T.R., Eveson J.P. 2019. Counting with DNA in metabarcoding studies: How should we convert sequence reads to dietary data? Mol. Ecol. 28:391–406.

Ficetola G.F., Pansu J., Bonin A., Coissac E., Giguet-Covex C., De Barba M., Gielly L., Lopes C.M., Boyer F., Pompanon F., Rayé G., Taberlet P. 2015. Replication levels, false presences and the estimation of the presence/absence from eDNA metabarcoding data. Mol. Ecol. Resour. 15:543–556.

Fofonoff P.W., Ruiz G.M., Steves B., Simkanin C., Jt &. C. 2018. *Lepomis cyanellus*. Available from https://invasions.si.edu/nemesis/species_summary/168132.

Frimpong E.A., Angermeier P.L. 2009. Fish traits: A database of ecological and life-history traits of freshwater fishes of the United States. Fisheries. 34:487–495.

Gotelli N.J., Colwell R.K. 2001. Quantifying biodiversity: procedures and pitfalls in the measurement and comparison of species richness. Ecol. Lett. 4:379–391.

Hammerson G. 2011. Dorosoma petenense. Available from https://explorer.natureserve.org/Taxon/ELEMENT_GLOBAL.2.105828/Dorosoma_petenense.

IDNR. 2024. Fish species - quillback. Available from https://programs.iowadnr.gov/lakemanagement/fishiowa/fishdetails/ULL.

iNaturalist. 2026a. Hydrophlox chiliticus. Available from https://www.inaturalist.org/taxa/1545658-Hydrophlox-chiliticus.

iNaturalist. 2026b. Greenhead shiner (Hydrophlox chlorocephalus). Available from https://www.inaturalist.org/taxa/1545657-Hydrophlox-chlorocephalus.

Lagacy E. 2024. Species Status Assessment: Swallowtail Shiner. Available from https://extapps.dec.ny.gov/fs/programs/dfw/SWAP2025/Freshwater.

Lamprecht S.D., Bettinger J., Scott M.C. 2015. Quillback Carpiodes cyprinus. Available from https://www.dnr.sc.gov/swap/supplemental/freshwaterfish/quillback2015.pdf.

LeGrand, H., J. Amoroso, and T. Howard. 2026. Freshwater fishes of North Carolina. Available from https://auth1.dpr.ncparks.gov/fish/index.php.

Macher T.-H., Schütz R., Arle J., Beermann A.J., Koschorreck J., Leese F. 2021. Beyond fish eDNA metabarcoding: Field replicates disproportionately improve the detection of stream associated vertebrate species. Metabarcoding Metagenom. 5.

Nico L., Fuller P., Neilson M., Daniel W., and Bartos A. 2025. Red Shiner (Cyprinella lutrensis) - Species Profile. Available from https://nas.er.usgs.gov/queries/factsheet.aspx?SpeciesID=518.

NJFW. 2026. Black Crappie (Pomoxis nigromaculatus). Available from https://dep.nj.gov/njfw/wp-content/uploads/njfw/Black-Crappie.pdf.

Oksanen J., Simpson G.L., Blanchet F.G., Kindt R., Legendre P., Minchin P.R., O'Hara R.B., Solymos P., Stevens M.H.H., Szoecs E., Wagner H., Barbour M., Bedward M., Bolker B., Borcard D., Carvalho G., Chirico M., De Caceres M., Durand S., Evangelista H.B.A., FitzJohn R., Friendly M., Furneaux B., Hannigan G., Hill M.O., Lahti L., McGlinn D., Ouellette M.-H., Ribeiro Cunha E., Smith T., Stier A., Ter Braak C.J.F., Weedon J. 2024. vegan: Community Ecology Package. .

Parker B.R., Franzin W.G. 1991. Reproductive biology of the quillback, Carpiodes cyprinus, in a small prairie river. Can. J. Zool. 69:2133–2139.

Pauly F.R.A. 2010. Fishbase. Available from https://www.fishbase.org/search.php.

Rohde F.C., Arndt R.G., Lindquist D.G. 1995. Freshwater fishes of the Carolinas, Virginia, Maryland, and Delaware. Chapel Hill, NC: University of North Carolina Press.

Shaw H.A.M.M. 2023. *Dorosoma cepedianum* Gizzard Shad. Available from https://explorer.natureserve.org/Taxon/ELEMENT_GLOBAL.2.104453/Dorosoma_cepedianum.

Shirazi S., Meyer R.S., Shapiro B. 2021. Revisiting the effect of PCR replication and sequencing depth on biodiversity metrics in environmental DNA metabarcoding. Ecol. Evol. 11:15766–15779.

Stauffer S., Jucker M., Keggin T., Marques V., Andrello M., Bessudo S., Cheutin M.-C., Borrero-Pérez G.H., Richards E., Dejean T., Hocdé R., Juhel J.-B., Ladino F., Letessier T.B., Loiseau N., Maire E., Mouillot D., Mutis Martinezguerra M., Manel S., Polanco Fernández A., Valentini A., Velez L., Albouy C., Pellissier L., Waldock C. 2021. How many replicates to accurately estimate fish biodiversity using environmental DNA on coral reefs? Ecol. Evol. 11:14630–14643.

Tracy B., Rohde F., Hogue G. 2020. An annotated atlas of the Freshwater Fishes of North Carolina. Southeast. Fishes Counc. Proc. 60.

U.S. Geological Survey. 2026. *Morone americana* (Gmelin, 1789). Available from https://nas.er.usgs.gov/queries/FactSheet.aspx?speciesID=777.

Wickham H. 2016. ggplot2: Elegant Graphics for Data Analysis. .

Wickham H. 2023. dplyr: a grammar of data manipulation. Available from https://user2014.r-project.org/abstracts/talks/45_Wickham.pdf.

Wilson T.M., Acre M.R., Williams F., Calfee R.D., Mayer C.M., Mapes R.L., Kemp C.M., Young R.T.,

Byrne M.E. 2025. Reproductive biology of invasive grass carp (Ctenopharyngodon idella) in two North American systems. Available from https://www.usgs.gov/publications/reproductive-biology-invasive-grass-carp-ctenopharyngodon-idella-two-north-american.

Zar J.H. 2014. Spearman rank correlation: Overview. Wiley StatsRef: Statistics Reference Online.
